# Notch-independent Her6 contributes to control of neural stem cell maintenance by shifting Notch signaling from lateral inhibition towards lateral induction mode

**DOI:** 10.64898/2026.09.16.752018

**Authors:** Jan Rothenpieler, Rebecca Veit, Dragana Stefanovska, Julia Schwab, Christian Sigloch, Dominic Grün, Wolfgang Driever

## Abstract

The growing brain faces the challenge to establish stable spatially defined neural proliferation zones that accomplish both neural stem cell (NSC) maintenance and dynamic neurogenesis. We investigated the relative impact of Notch signaling, Notch-dependent, and Notch-independent HES/Her transcription factors on target gene expression in the larval zebrafish brain. Both genetic evidence and transcriptional network analysis reveal a pivotal role of Her6, a Notch-independent HES1 homolog, in NSC control. Her6 at G1, and Notch signaling at G2, regulate distinct cell cycle control genes, favoring arrest. Her6 and Notch activity combined differentially control *delta*, *jagged*, *lfng* and *notch3* expression to potentially shift Notch signaling mechanisms from Delta-driven lateral inhibition to Jagged-Notch3-mediated lateral induction. Indeed, we detect *in situ* spatial expression profiles consistent with lateral induction in high *her6* expressing regions of the larval and adult brain. Her6-dependent lateral induction may spatially organize coherent populations of long-term NSCs in active proliferation zones.

## Introduction

During development of the vertebrate nervous system, the constraints governing stability, dynamics, and plasticity of neural stem (NSC) and progenitor cell (NPC) populations change dramatically. These processes are prominently regulated by Delta-Notch signaling and its targets, including Hairy and Enhancer-of-split related (HES/Her) transcriptional repressors (Artavanis-Tsakonas et al., 1999; Kageyama et al., 2007), which together control NSC maintenance, the balance between quiescence and activation of NSCs (Chapouton et al., 2010; Engler et al., 2018; Sueda et al., 2019; Sood et al., 2022; Lampada and Taylor, 2023), as well as neurogenesis progression (Hitoshi, 2002; Hatakeyama et al., 2004). In the early embryonic neuroectoderm, stemness is maintained and neurogenesis inhibited by constitutive expression of HES/Her repressors in a Notch signaling independent manner, including Her3 in zebrafish (Onichtchouk et al., 2010) and Hes3 in mice (Katoh and Katoh, 2007). Subsequently, in the neural tube lateral inhibition by Delta-Notch signaling controls dynamic neurogenesis (Chitnis et al., 1995; Artavanis-Tsakonas et al., 1999). The major growth of the brain at late embryonic and larval stages is preceded by establishment of highly active zones, termed neural proliferation zones (NPZs) in zebrafish, that ensure both, stable NSC populations and strong neurogenesis (Wullimann and Knipp, 2000; Kaslin et al., 2008). For mice, a concept has been developed that in “boundary” regions, like the mid-hindbrain boundary, floor plate, or zona limitans intrathalamica (ZLI), high Hes1 expression maintains NSCs, while in adjacent “compartments” oscillating Hes1 and Hes5 control neurogenesis progression (Baek et al., 2006; Kageyama et al., 2007). Whether Hes1 and Hes5 may have fundamentally different activities in boundaries and compartments is not well understood.

In the *Drosophila* embryo, *hairy* represses neurogenesis in the neurogenic ectoderm in a Notch-independent (NI) manner during early patterning (Orenic et al., 1993). In contrast, after establishment of neurogenic stem cell zones, Notch-dependent (ND) expression of *Enhancer of split* mediates lateral inhibition in proneural clusters during neuroblast segregation (Artavanis-Tsakonas et al., 1999). HES/Her transcription factors in vertebrates are similarly regulated by ND and NI mechanisms. In mouse, Hes1 and Hes5 are Notch effectors, though Hes1 is also regulated by NI input (de la Pompa et al., 1997; Ohtsuka et al., 1999; Kageyama et al., 2007; Riya et al., 2026). In zebrafish, a larger family of HES/Her factors controls neural development (Geling et al., 2004; Bae et al., 2005; Chapouton et al., 2011; Sigloch et al., 2023). While the zebrafish Hes5 homologs (*her2*, *her4.1*-*4.5*, *her12*, *her15.1-15.2*) are ND, expression of the Hes1 homologs *her6* and *her9* is largely NI (Bae et al., 2005; Chapouton et al., 2011; Sigloch et al., 2023). Similar to Hes1, Her6 shows higher oscillating expression in “boundary” regions like ZLI and floor plate, and lower oscillating expression in active neurogenesis compartments (Soto et al., 2020; Sigloch et al., 2026). Combined NI *her* gene mutations cause more severe NSC deficiency than combined ND *her* mutations, and, surprisingly, a single wildtype *her6* allele can compensate for loss of other ND and NI *her* genes in compound mutants (Sigloch et al., 2023). The mechanisms behind this unique role of Her6 in NSCs, and more general the differences between ND and NI *Hes/her* gene activities, are not well understood.

There is accumulating evidence that distinct Notch signaling modes regulate non-cycling NSCs (ncNSCs) and proliferating NSC (pNSC) populations, as well as NPC lineage progression. Notch1 signaling in both mice and fish prevails in pNSCs and NPCs to inhibit neuronal differentiation (Basak et al., 2012), while Notch3 (and also Notch2 in mice) signaling is required to maintain ncNSCs in their quiescent state (Alunni et al., 2013; Kawai et al., 2017; Engler et al., 2018; Than-Trong et al., 2018). Although Notch1, Notch2 and Notch3 can be activated by both Delta and Jagged, differences in their binding affinities, signal activation and spatial distribution give rise to distinct Notch signaling outputs. Glycosylation by Lunatic fringe (Lfng) enhances Notch1 activation by Delta but not by Jagged, while Jagged induces stronger activation of Notch2 and Notch3 in the absence of Lfng (Shimizu et al., 1999; Kakuda and Haltiwanger, 2017; Kuintzle et al., 2025; Ortica et al., 2026). Delta-Notch1 signaling has been shown to control neurogenesis progression throughout vertebrates (Chitnis et al., 1995), while Jagged-Notch2/Notch3 signaling has been demonstrated to be required for NSC maintenance in mice (Nyfeler et al., 2005; Lavado and Oliver, 2014) and zebrafish (Than-Trong et al., 2018; Ortica et al., 2026). Jagged and Delta signaling have been suggested to have different patterning outcomes: Delta-Notch1 through lateral inhibition generates salt-and-pepper patterns of NSCs and NPCs, while Jagged-Notch3 through lateral induction may promote coherent zones of NSCs (Bocci et al., 2020). Although Jagged and Delta signaling are clearly involved in regulating distinct NSC and NPC states, it is unclear how the transition between the two signaling states is regulated, and how both together contribute to spatial patterning of NPZs.

Here we analyze larval zebrafish NSC and NPC transcriptomes to reveal the molecular diversity and dynamics of ND and NI *her* genes expressing neural populations. To elucidate distinct and shared regulatory networks downstream of Notch signaling, we employ heat-shock induced Her4, Her6 and NICD overexpression followed by time-series transcriptome analyses. We identify ncNSCs already at 3 days post fertilization (dpf), and find that both Her6 and NICD through distinct transcriptional targets may promote cell cycle arrest or exit, and quiescence. Surprisingly, ND Her4 downstream transcriptional repression is largely limited to ND *her* genes. In contrast, Her6 and NICD control shared repression of proneural and *delta* genes, but distinct regulation of Notch signaling pathway components, including *notch3, jag1b,* and *lfng*. We propose a novel role for Her6 in NSCs by shifting Notch signaling mechanisms from Delta-Notch1-mediated lateral inhibition towards Jagged-Notch3-mediated lateral induction. We find patterns of gene expression in the adult zebrafish telencephalic ventricular proliferation zone that support this hypothesis.

## Results

### *her6* expression in non-cycling and proliferating NSCs

To better characterize *her6* expression and function along NSC and NPC lineages, we FACS-isolated Her6-mNeonGreen (Her6-mNG) expressing cells to perform both singe-cell (sc) and bulk RNA-seq (Figs. 1 and S1). We used a CRISPR/Cas9 knock-in of mNeonGreen (mNG) fluorescent protein in frame at the carboxy-terminus of the *her6* coding region that faithfully reports Her6 expression (Sigloch et al., 2026), focusing on 3 dpf zebrafish larvae, when NPZs are established. Given that high versus oscillating low expression modes of Hes1 were reported for distinct NSC populations in mice (Baek et al., 2006; Ochi et al., 2020), and distinct zebrafish Her6 expression levels have also been observed (Sigloch et al., 2026), we aimed to determine whether different levels of Her6 would correlate with distinct neurogenesis states. Therefore, we separately collected cells with low and with high Her6-mNG fluorescence (Fig. 1A). *her6* and *mNG* read counts were indeed higher in high Her6-mNG fluorescence bulk RNA-seq samples (Fig. S1A,B). Both high and low mNG-fluorescent cells are enriched for expression of a large shared set of genes linked to NSCs and radial glia (RG). Further, expression of quiescence (*fabp7a*, *glula*, *ptn*, *mdka*, *sox3*), but also proliferation markers (*mcm2*, *mcm4*, *mcm5*, *mcm6*, *plk1*; Figs. 1B and S1C,E) was detected, while Gene Ontology (GO) analysis revealed depletion of genes associated with neuronal differentiation (Fig. S1D). Interestingly, while the Notch downstream targets *her4.1*, *hey1 and hey2* are enriched in Her6-mNG positive cells, neither *delta* nor *notch* genes are differentially expressed. In contrast, we found the Notch ligand *jag1b* to be enriched in Her6-mNG positive cells (FC 2.3, *p* 0.003). Together these data show that Her6 is primarily expressed in NSCs, along with direct Notch downstream targets, and that quiescence markers are already detected at 3 dpf.

**Figure 1.**
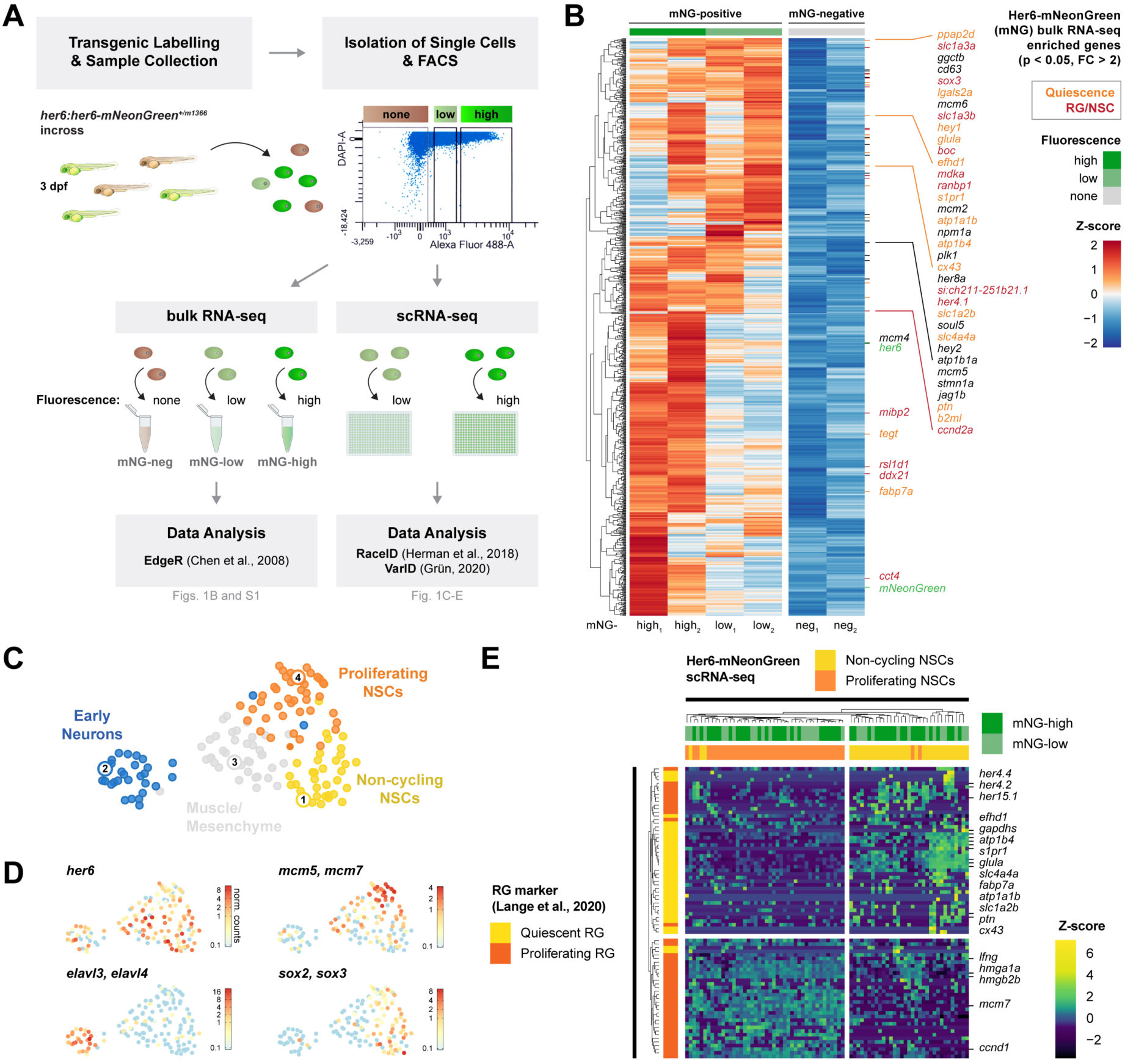
NI *her6* is expressed in non-cycling and proliferating NSCs along with Notch downstream targets. **(A)** Schematic workflow for bulk and scRNA-seq of FACS-purified Her6-mNG cells from 3 dpf zebrafish larvae. **(B)** Heatmap of 905 genes enriched in Her6-mNG positive (high or low) samples. Selected genes are shown. Marker genes for quiescence (orange) and RG/NSC (red) are highlighted. Depleted genes are depicted in **Fig. S1E**. Differences between high and low mNG samples may in part be explained by high mNG samples having a higher contribution of boundary NSCs with a tendency to express diverse region-specific markers. **(C)** UMAP representation of *VarID* clustering of 146 high-quality cells (see Methods) from Her6-mNG scRNA-seq analysis revealed four clusters with 34 (cluster mNG#1), 31 (mNG#2), 38 (mNG#3), and 43 (mNG#4) cells. **(D)** Log2 normalized expression of selected genes in clustering from **C**. **(E)** Hierarchical clustering of all cells from NSC clusters mNG#1 and mNG#4 from **C** using a published set of adult-type qRG and pRG markers (Lange et al., 2020). Selected genes are shown.

Our corresponding scRNA-seq analysis distinguishes ncNSCs and pNSC clusters (Fig. 1C,D). A heatmap visualizing expression of quiescence and proliferating RG markers (Lange et al., 2020) confirms this cluster assignment (Fig. 1E), and reveals that cells with high or low mNG fluorescence locate to both clusters (Fig. 1E, green cell annotation). However, we observed that cells with higher *her6* transcript counts tend to map to the ncNSC cluster (*her6* was 2.85-fold enriched in ncNSCs vs. pNSCs; *p* 5.969e-08; Fig. 1D). Thus, *her6* oscillation may cause variable *her6* mRNA read counts in ncNSC cluster cells, but *her6* levels are typically lower when cells progress into pNSCs.

### ncNSCs with distinct *hes/her* expression profiles are already established in the 3 dpf larval brain

To better understand the activities of *her* genes and Notch signaling in NSCs, NPCs, and early neurons, as well as during lineage transitions (Fig. 2A), we expanded our analyses to published scRNA-seq data of 3 dpf larval zebrafish heads (Raj et al., 2020). We extracted a subset of cells expressing, alone or in combinations, SoxB1 members as markers for NSCs and NPCs (*sox2, sox3*), and the following neural *hes/her* genes: (1) Hes1 homologs NI *her6* or *her9*, and (2) Hes5 homologs ND *her4.1-4.5, her12*, *her2,* or *her15.1-15.2*, as well as (3) Hes6 homologs ND *her8a*, *her8.2,* or NI *hes6* (Sigloch et al., 2023; Chen et al., 2024). Together, these cells define our “HER/SOX-3dpf” dataset (see Methods, Fig. 2B). Using *VarID* for cluster analysis we identified nine clusters correlating to distinct neurogenesis stages (Fig. 2C-G): There are two NSC clusters (*sox2*, *sox3*, *id1*, *mdka*), one with higher levels of quiescence markers (#14 *cx43, ptn, glula*) and one representing pNSCs (#15; *mki67*, *mcm7*, *cdk1*). Three progenitor clusters (#1, 8 and 9) express proneural genes (*neurog1*, *ascl1a*, *ascl1b*) and proliferation markers (*pcna*, *mki67*), with #8 and 9 expressing also progressively late NPC markers (*neurod4, elavl3*). Four clusters represent early neurons (#2, 4, 6 and 7; *elavl3*, *elavl4*). Non-neural and mature neuron clusters were not considered in further analyses (grey clusters in Figs. 2C and S2, see Methods). To identify potential subtypes, we re-clustered cells of the NSC (#14 and 15) and the NPC (#1, 8 and 9) clusters and analyzed cell cycle, *her* gene, and Notch pathway component expression (Fig. S3 and Methods). This confirmed distinct subclusters of ncNSCs and pNSCs (Figs. 2D and S3C,D).

**Figure 2.**
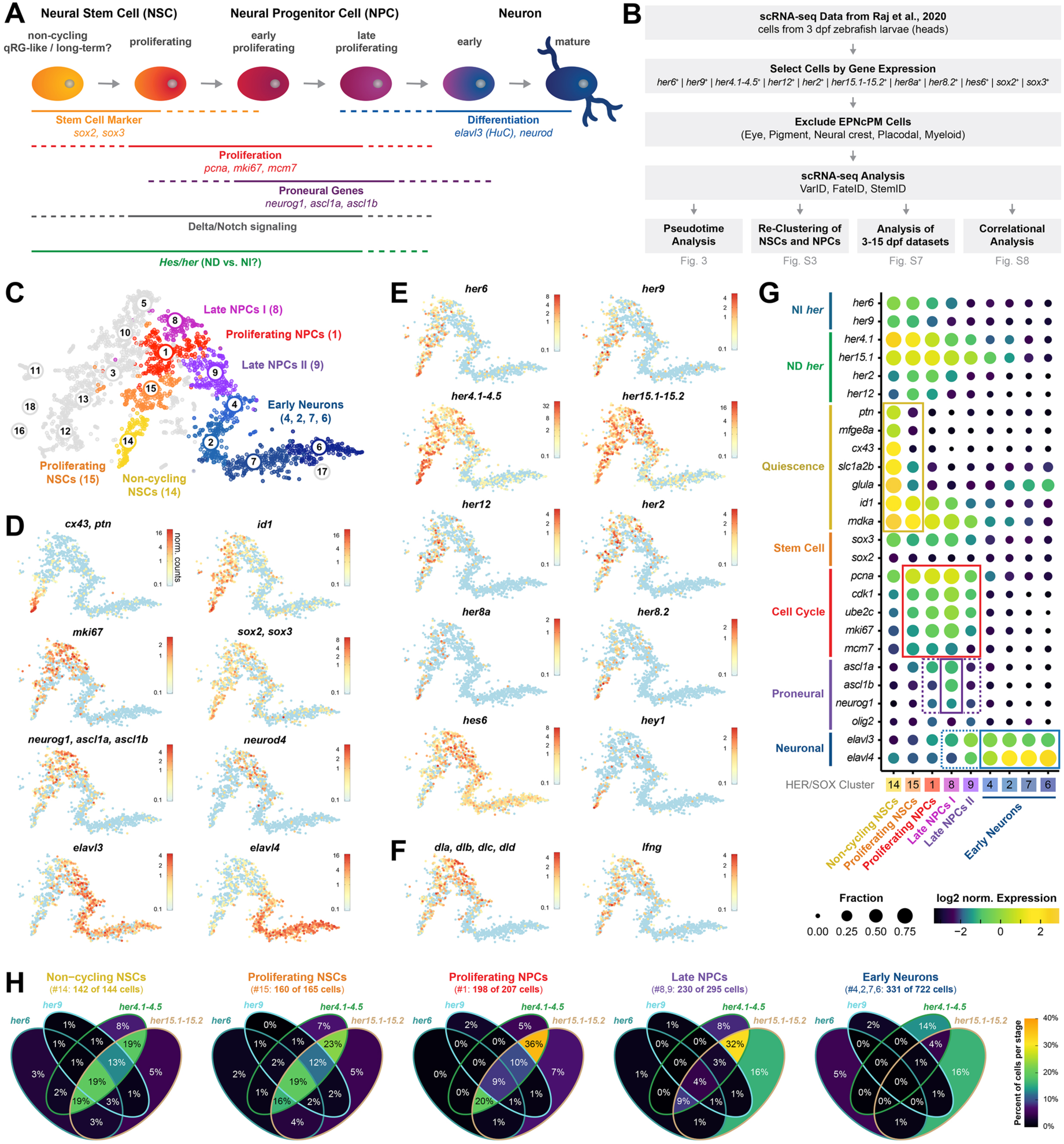
NI and ND *her* genes are co-expressed in NSCs, but in NPCs ND *her* genes prevail. **(A)** Scheme of cell lineage from ncNSC to neuron with examples of marker gene expression profiles during neurogenesis. **(B)** Selection of the HER/SOX dataset (see Methods) and schematic workflow for analyses of publicly available scRNA-seq data (Raj et al., 2020). **(C)** tSNE map of *VarID* clustering from scRNA-seq analysis of 2555 high-quality cells reveals 9 neural lineage clusters with 144 (ncNSCs, #14), 165 (pNSCs, #15), 207 (pNPCs, #1), 108 and 187 (late NPCs, #8 and 9), and 116, 172, 242 and 192 cells (early neurons, clusters #4, 2, 7 and 6). Non-neural and outlier clusters (grey) were not considered in further analyses (**Fig. S2B**). **(D-F)** Log2 normalized expression of neurogenesis markers (**D**), *her/hes/hey* genes (**E**) and Notch signaling pathway components (**F**) in neural clusters from **C**. **(G)** Fraction-dotplot with the expression of depicted marker genes in neural clusters from **C**. **(H)** Proportion of cells per neurogenesis stage with combined and single *her* gene expression. Only the depicted *her* genes are considered.

*her6, her9*, *her4* and *her15* genes are all highly expressed in ncNSCs (Fig. 2E,G), with *her15* genes also expressed at high levels in late cycling NSCs and all NPCs, in which *her2* and *her12* are also expressed. *delta* genes and *lfng* are expressed predominantly in pNSCs (#15) and NPCs, but in comparison rather low in ncNSCs (Figs. 2F and S4A-C). ncNSCs also express RG, astrocyte, and other NSC markers, while cell cycle genes are silent. In comparison to adult RGs (Lange et al., 2020), the ncNSCs express genes enriched in *her4.1*+ quiescent RGs (qRG) but not in proliferating RGs (pRG). A recent scRNA-seq study (Morizet et al., 2024) identified multiple qRG clusters with heterogenous quiescence depth in the adult zebrafish telencephalon, and defined a gene expression profile characteristic for qRGs. Many of these qRG-defining genes, including *ptn* and *mfge8a*, are also expressed in our 3 dpf ncNSCs, which appear as a rather homogenous cluster (Fig. S3). The dataset from Morizet et al. (Morizet et al., 2024) also revealed expression of NI *her6* and *her9*, and of ND *her4.1* and *her15.1* in adult qRG clusters (Fig. S5A,D). In summary, we find that 3 dpf larval ncNSCs already have a transcriptional signature common with adult qRGs.

### Dynamic expression profiles of NI and ND *her* genes in neural stem and progenitor cells

To investigate the expression dynamics of NI and ND *her* genes during neurogenesis progression at single-cell level, we performed pseudotime analysis based on the HER/SOX-3dpf cell clusters, using *VarID* (Gruen, 2020) and *FateID* (Herman et al., 2018) (Fig. 3A,B and Methods). We found that trajectory inference accurately reflects neurogenesis stages (Fig. 3C), with cells expressing markers for stemness, proliferation or differentiation being ordered along pseudotime (Fig. 3D). A self-organizing map of the pseudo-temporal expression profiles grouped similar expression profiles of genes into modules (Figs. 3E, S6). Plotting of pseudo-temporal expression of individual genes (Fig. 3F) enabled clear discrimination of early (G1/S) and late (S/G2) cell cycle phases. Markers for quiescence are strongly expressed in cells of the ncNSC clusters, and rapidly and completely downregulated when cells enter the cell cycle. NI *her6* and *her9* are expressed at highest levels in ncNSCs and are downregulated during cell cycle progression. Similarly, ND *her4* genes are highly expressed in NSCs, and remain expressed into NPC stages. In contrast, ND *her15* genes are expressed at lower levels in ncNSCs and reveal elevated expression in pNSCs and NPCs until differentiation, when *elavl4*, *neurod6a*, and *neurod6b* are expressed. The proneural genes *neurog1*, *ascl1a* and *ascl1b* are expressed in cycling cells. Surprisingly, expression of *delta* genes, *lfng* and *notch1a* is low in ncNSCs, but peaks along with late cell cycle genes in NPCs, when also *elavl3, neurod1* and *neurod4* start to be expressed.

**Figure 3.**
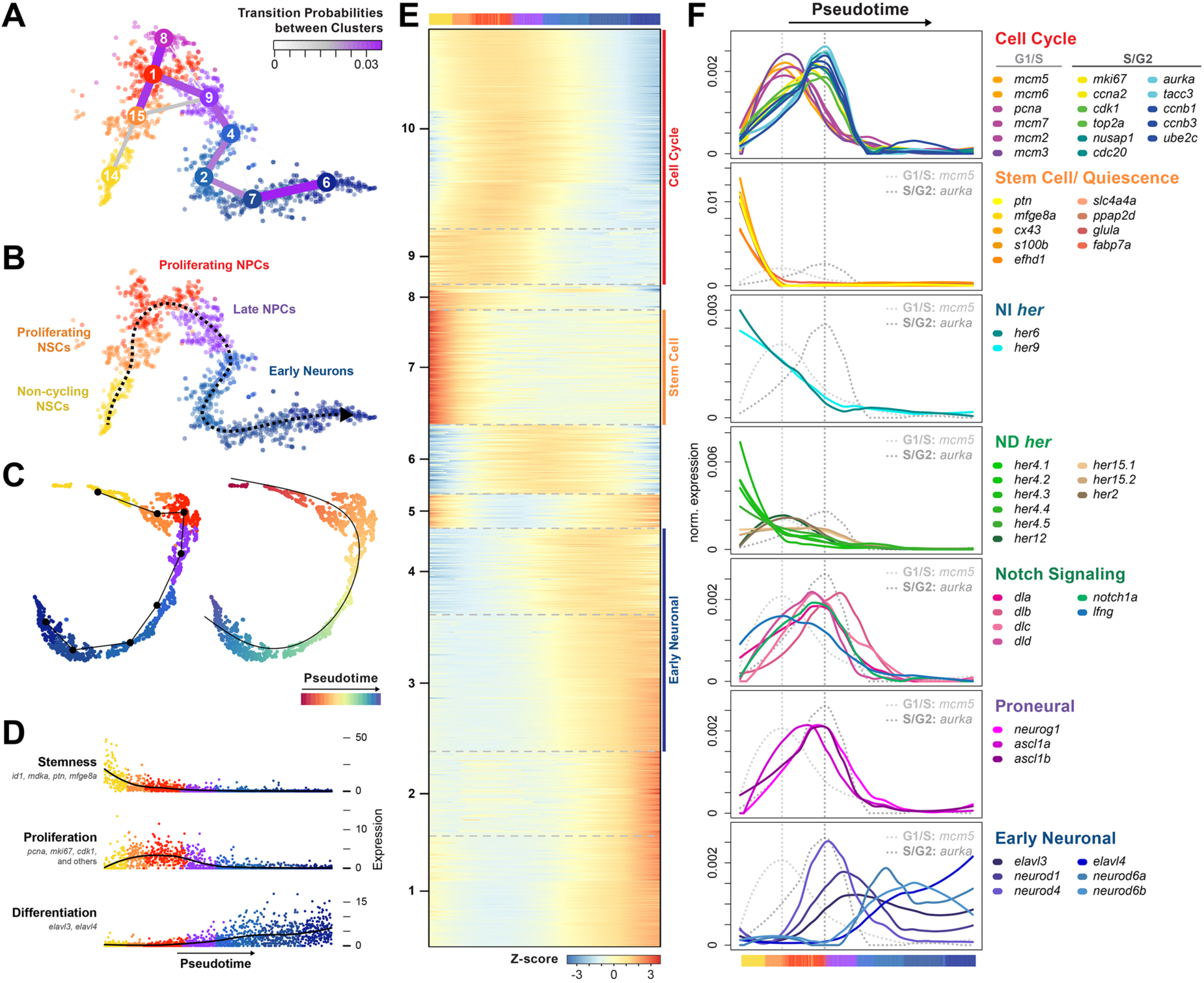
Pseudotime analysis reveals correlation of NI and ND *her* expression profiles with markers for quiescence and progression of neurogenesis. **(A)** tSNE map with transition probabilities between clusters based on Fig. 2C. **(B)** Lineage included in pseudotime. **(C)** UMAP representation of clusters #14, 15, 1, 9, 4, 2, 7 and 6 from **A**, with the continuous trajectory indicated (**left**) and pseudotime highlighted (**right**). **(D)** Pseudo-temporal expression profiles of genes for stemness, proliferation and neuronal differentiation, respectively. Additional included proliferation markers: *ccna2*, *ccnb1*, *ccnb2*, *ccnb3*, *ccnd1*, *ccnd3*, *ccne1*, *ccne2*, *cdt1*, *mcm5*, *mcm7*, *nus1*, *skp2*, *tmem2,* and *ube2c*. **(E)** Heatmap with Z-score pseudo-temporal expression profiles (rows) for cells in pseudo-temporal order (columns), derived from self-organizing map (SOM). Colors (top) indicate cluster affiliation of cells (color code as in **A**). The SOM modules are indicated on the left. Selected genes are highlighted in **Fig. S6**. **(F)** Loess-smoothed pseudo-temporal expression profiles of selected genes. Cluster affiliation of cells is indicated at the bottom by colors as in **A**.

Given that ncNSC populations have previously not been observed as early as 3 dpf in larvae, we asked whether the observed expression profiles of NI *her* and Notch signaling pathway components may persist through juvenile stages. We selected HER/SOX cells from 3, 5, 8, and 15 dpf scRNA-seq data (Raj et al., 2020) and clustered the dataset integrating all stages (Fig. S7A,B,F). The analysis revealed that ncNSCs (cluster 21) of all stages are characterized by similar cell cycle, stemness and neurogenesis markers (Fig. S7C,D). Further, the analysis of *her* and *delta* genes and of *lfng* expression across all stages indicates that the distinct NSC and NPC clusters show similar expression profiles of *her* and Notch signaling genes across all developmental stages (Fig. S7E). Thus, dynamic changes of NI and ND *her* expression and Notch signaling pathway genes during lineage progression from ncNSCs to early neurons appears to be conserved across developmental stages.

We next aimed to better characterize the combined activities of ND and NI Her in individual NSCs and NPCs, focusing on Hes1 and Hes5 homologs (Figs. 2H and S4D-F). We found that more than 80% of NI *her6* and/or *her9* expressing cells also express ND *her* genes, while almost two thirds of ND *her* positive cells do not have transcript counts for NI *her* genes (Fig. S4E). Across all clusters, few individual cells express only single *her/hes* genes, except *hes6* in late NPCs and early neurons (Fig. S4F bottom). Combined expression of NI and ND *her* genes is characteristic for ncNSCs (62% of cells) and pNSCs (59%; Figs. 2H and S4F). In pNPCs the balance shifts from NI and ND *her* genes combined (42%) to ND *her* genes only (48%). Late NPCs predominantly express exclusively ND *her* genes (56%) while only 19% express a combination of NI and ND *her* genes.

The simultaneous expression of NI and ND *her* genes in NSCs may indicate that distinct regulatory activities of both classes of *her* genes together are required for specific NSC properties. In addition, since Her6 expression oscillates (Soto et al., 2020; Doostdar et al., 2024; Sigloch et al., 2026), high versus low level *her6* expressing cells may correlate with distinct transcriptional profiles and potentially NSC states. We investigated these questions by identifying genes expressed at levels correlated or anticorrelated to *her* gene expression levels (Fig. S8; for pseudo-bulking see Methods). Both *her6* and *her4.1* show highly correlated expression with multiple stem cell and quiescence markers (Fig. S8B). For *her6, her4.1* and *her15.1* separately, we mapped the ranked sum of expression of the top 50 correlated genes onto the HER/SOX-3dpf NSC and NPC clusters (Fig. S8C). The 50 genes with expression levels highly correlated to *her6,* or respectively to *her4.1*, in bulk map to the lineage states, specifically highest levels to ncNSCs, and intermediate levels to pNSCs, but not to the respective *her* expression levels in individual cells. Such a strict correlation to lineage states is not observed for *her15.1*. We conclude that instantaneous high versus low *her6* and *her4.1* expression does not specify distinct temporally dynamic oscillating NSC transcriptional states, but oscillating *her* expression levels at the target transcriptome may be integrated over time, and change only during lineage progression from ncNSC to pNSC states.

### Identification of Her6, Her4 and NICD transcriptional targets by induced overexpression and time-series transcriptome analyses

Our data and genetic evidence (Sigloch et al., 2023) suggest that NI and ND *her* genes make distinct contributions to control of NSCs, NPCs, and lineage progression. Thus Her6 may have distinct transcriptional targets as compared to Notch signaling and Her4. To uncover downstream targets of Her6 and Her4, we used heat-shock-driven overexpression (OE) of *her6-FLAG* (Sigloch et al., 2023) (here “Her6-OE”) and *her4.1-FLAG* (Sigloch et al., 2023) (here “Her4-OE”). Further, to identify Notch signaling transcriptional targets more broadly, we used heat-shock Gal4 driven overexpression of *notch1a-ICD* (Scheer and Campos-Ortega, 1999) (here “NICD-OE”). We heat-shocked larvae *in vivo* at 2 dpf, and sampled larval brains before onset of heat-shock and at eight time points between 0.5 and 6 h post heat-shock (pHS), and performed bulk-RNA-seq (Fig. 4A). To be able to distinguish early, potentially direct, from later indirect transcriptional responses, we sampled at 30 min intervals in the first 3 h.

**Figure 4.**
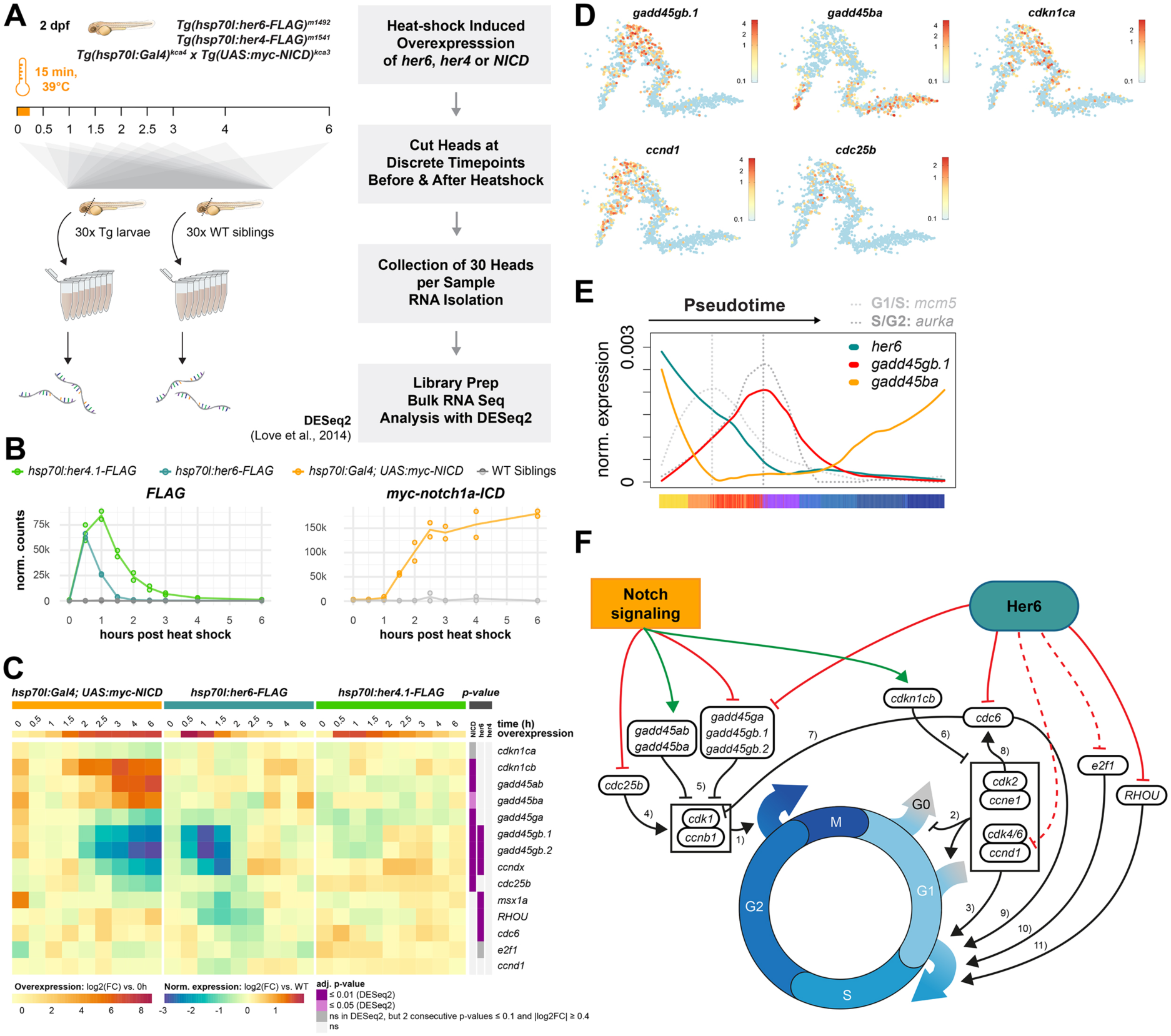
Time series analyses of *her4.1*, *her6* or *NICD* overexpression reveal distinct and shared transcriptional targets involved in cell cycle control. **(A)** Schematic workflow of bulk RNA-seq time series before and after NICD-OE, Her6-OE or Her4-OE. **(B)** Expression of *her4.1-FLAG*, *her6-FLAG* and *myc-NICD* transgenes in OE experiments. **(C)** Heatmap with log2(FC) expression of selected cell cycle genes in OE experiments. **(D)** Expression of selected cell cycle genes in HER/SOX-3dpf dataset. **(E)** Opposing pseudo-temporal expression profiles of *gadd45gb.1* and *gadd45ba* (compare Fig. 3F). **(F)** Schematic diagram of regulatory impact of Notch signaling (NICD-OE) and Her6 activity on expression of cell cycle regulators. While Notch signaling predominantly regulates G2/M progression, Her6 mainly acts on the G1/S transition. The downstream effects (black connectors) of the affected factors have been previously published: (1-3) (Malumbres, 2014; Pellarin et al., 2025), (4) (Lammer et al., 1998), (5) (Vairapandi et al., 2002), (6) (Lee et al., 1995; Matsuoka et al., 1995), (7) (Clay-Farrace et al., 2003; Lau et al., 2006), (8) (Mailand and Diffley, 2005), (9) (Liang and Stillman, 1997), (10) (Muller et al., 1997), (11) (Tao et al., 2001).

Experimental setup and data analysis aimed at eliminating heat-shock and Gal4 effects on transcription from processed data (Methods, Figs. S9 and S10). *her6*-*FLAG* transcripts display a quick increase that peaks at 0.5 h pHS and decays to less than one tenth of the peak level within 1 h, while *her4.1*-*FLAG* transcripts peak at 1 h pHS and fall below one tenth of the peak within 2 h (Fig. 4B, C). For *notch1a* we observed increased reads from 1.5 h pHS on, saturating at 2.5 h and persistent to 6 h pHS (Fig. 4B, C) revealing the heat-shock Gal4 to cause a slower but sustained increase in NICD expression, due to delay in the dual expression system and the stability of the Gal4 protein (Scheer et al., 2002).

### Her6 and NICD differentially regulate genes involved in cell cycle control

Given that Her6 prevails in ncNSCs, while Notch signaling is also active in pNSCs and NPCs, we asked whether Her6- and NICD-OE may differentially affect cell cycle control genes (Figs. 4 and S11). We found that both, Her6- and NICD-OE, most severely affected genes of the *gadd45* gene family, genotoxicity-induced survival factors which inhibit the Cdk1/CyclinB complex, leading to G2/M cell cycle arrest and preventing apoptosis (Vairapandi et al., 2002; Gupta et al., 2006). *gadd45ab* and *gadd45ba* are both upregulated upon NICD-OE, while *gadd45g* paralogs were repressed by both Her6*-* and NICD-OE (Fig. 4C). Our pseudotime analysis revealed that these *gadd45g* paralogs are mainly expressed in late cycling NPCs, where *her6* is only sparsely expressed, while *gadd45ba* is most abundant in quiescent and differentiating cells, where the cell cycle is arrested (Figs. 4D,E and S6). Further, NICD-OE repressed *cdc25b*, which is essential for mitotic entry by dephosphorylation of Cdk1 (Kumagai and Dunphy, 1991). When checking for CDK inhibitors, we found Notch activity upregulated the *p57^kip2^* homolog *cdkn1cb*, while its paralog *cdkn1ca* appeared to be slightly repressed.

In addition, we found Her6 to repress *RHOU* (*si:ch211-133l11.10*), *cdc6, msx1,* and *ccndx.* We also observed a slight reduction of *ccnd1* and *e2f1* expression at two time points. The Rho GTPase RHOU is a Wnt responsive Cdc42 homolog, and known to promote cell cycle entry from quiescence in mouse cells (Tao et al., 2001). Cdc42 has been shown to activate *Ccnd1* transcription in cell culture (Bauerfeld et al., 2001). Therefore, as *cdc42* is unaffected upon Her6-OE (Fig. S11), downregulation of *ccnd1* could be indirect through RHOU regulation. Further, Her6-OE repressed *cdc6*, which is expressed in proliferating cells and part of the pre-replication complex required for G1/S transition (Yan et al., 1998; Lau et al., 2006), and is also involved in G2/M checkpoint activation by *cdk1* inhibition (Clay-Farrace et al., 2003; Lau et al., 2006). Additionally, *msx1a* is suppressed by Her6-OE. In *Drosophila msh/msx*has been implicated in the induction of *p57^kip2^* (the ortholog of *cdkn1ca*) (Otsuki and Brand, 2019), and in mice, Hes1 has been shown to repress *p57^kip2^* (Georgia et al., 2006; Monahan et al., 2009). Finally, the main driver for S-phase entry *e2f1* appeared to be slightly repressed (Fig. S11B).

In summary, the observed effects are consistent with a model in which Her6 predominantly inhibits cell cycle progression from G1 to S phase and to a lesser extend reentry into the cell cycle from G0, while Notch signaling regulates the cell cycle in a more complex fashion, including via inhibiting progression from G2 into M phase, potentially regulating a G2 arrest state (Fig. 4F).

Interestingly, we also found both *insm1* paralogs to be repressed by Her6- and NICD-OE (Fig. 5A). In mouse, Gadd45g and Insm1 were previously shown to be direct targets of *Hes1* (Kobayashi et al., 2009), and to promote conversion of apical to basal RG, and NPC delamination by repressing apical adherence junction proteins (Tavano et al., 2018). In zebrafish, *insm1a* and *insm1b* were reported to be expressed in the nervous system in patterns similar to proneural markers and *elavl3*, reflecting progenitor exit into differentiation (Lukowski et al., 2006). Our findings suggest that in Her6-positive NSCs Insm1 is downregulated to prevent delamination and premature exit from the stem cell niche, consistent with a role of Her6 in NSC maintenance.

**Figure 5.**
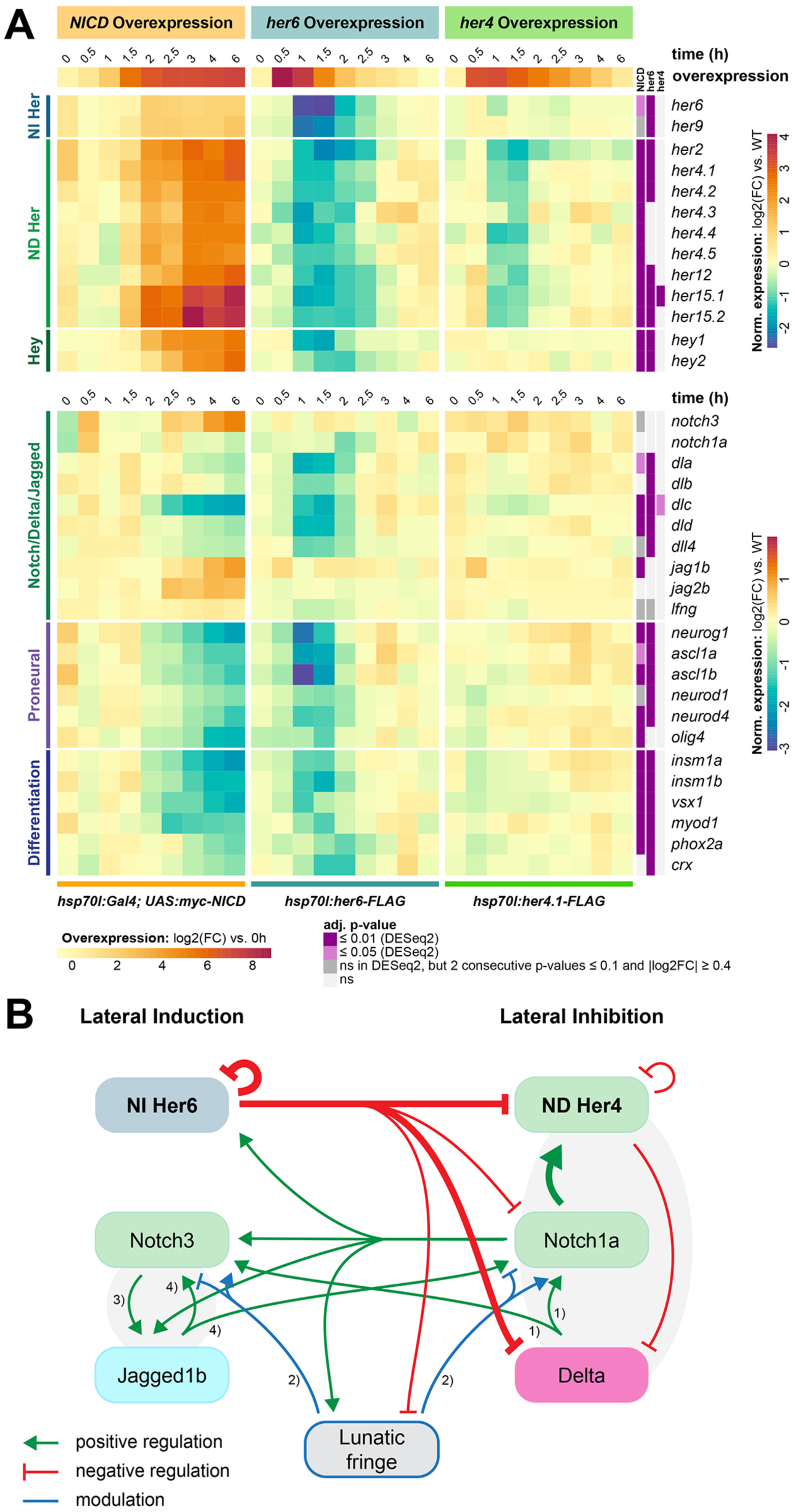
Time series analyses of *her4.1*, *her6* or *NICD* overexpression reveal distinct and shared downstream transcriptional targets regulating NSCs and neurogenesis. **(A)** Time series heatmap with log2(FC) expression of depicted target genes in OE experiments. Expression of additional genes is visualized in **Figs. S10-S13**. **(B)** The regulatory interactions identified in this study were used to develop a model revealing the potential of Her6 to shift Notch signaling from lateral inhibition and lateral induction modes. Molecular interaction not shown in this study but crucial for the model are based on the following literature: (1) (Jarriault et al., 1998; Shimizu et al., 2000), (2) (Kakuda and Haltiwanger, 2017; Kuintzle et al., 2025), (3) (Choi et al., 2008), (4) (Lindsell et al., 1995; Shimizu et al., 2000).

### NI Her6 and NICD differentially regulate Notch signaling pathway components

Our overexpression experiments revealed strong regulation of Notch pathway and neurogenesis genes (Figs. 5A, S12 and S13A,D), while no consistent pattern of regulation of WNT- or BMP-signaling pathway components was observed (Fig. S13B,C). Her6-OE resulted in significant downregulation of most Hes/Her/Hey-family genes (Figs. 5A and S12A,B) as well as of proneural genes including *ascl1a*, *ascl1b* and *neurog1* (Figs. 5A and S12A-D), in line with previous studies of *her6* in zebrafish (Scholpp et al., 2009; Sigloch et al., 2023) and Hes1 in mouse (Kageyama et al., 2000). Our data further revealed repression of all *delta* genes (*dla*, *dlb*, *dlc*, *dld*, *dll4*) by Her6, whereas *jagged* genes (*jag1a*, *jag1b*, *jag2a*, *jag2b*) were not affected (Figs. 5A and S13A). In contrast, Hes1 has been reported to directly repress *Jagged1*, together with *Dll1* and *Dll4,* in mouse (Kobayashi et al., 2009; Zhang et al., 2021). Interestingly, *lfng* and also *notch1a* were slightly repressed by Her6, whereas the expression of other Notch receptors was not notably affected. Finally, we found Her6 to repress genes involved in differentiation (including *vsx1*, *phox2a,* and *crx*; Figs. 5A and S10B). When we compared our findings to previously identified Hes1 ChIP targets in mouse (Kobayashi et al., 2009), we found that 14 out of 53 described Hes1 ChIP targets have paralogs or paralog groups regulated by Her6 in zebrafish, suggesting that significant components of the Hes1 / Her6 target gene network are conserved.

Her4-OE only affected expression of a very small number of genes (Fig. S10C). While we observed downregulation of all ND *her* genes, NI *her6* and *her9* were not affected (Figs. 5A and S12A,B). Further, only *dlc* expression was mildly repressed upon Her4-OE, while a significant regulation of other Notch ligands and proneural genes was not detected. These findings suggest that NI Her6 may control both NSC states and Notch signaling, while ND Her4 activity may be more restricted to modulation of Notch signaling activity.

NICD-OE resulted in upregulation of most Hes/Hey family genes (Figs. 5A and S12A,B), except for delayed downregulation (4 h pHS) of both *hes2* paralogs, which may have been downregulated indirectly through other *her* genes. *dlc* was strongly repressed upon NICD-OE, while the other *delta* genes were less repressed, especially when compared to their repression by Her6-OE (Figs. 5A and S13A, D). The Notch modulator *lfng* was slightly upregulated upon NICD-OE, which is in contrast to the slight repression by Her6-OE (Fig. 5A). We found *jag1b* expression to be significantly upregulated upon NICD-OE, and also observed a slight upregulation of *jag2b*. Finally, NICD-OE also upregulated *notch3*, while expression of other *notch* genes was unaffected (Figs. 5A and S13A,D).

Based on the interactions identified in our NICD- and Her6-OE experiments, we built a regulatory network model (Fig. 5B) in which Her6 represses Delta signaling as well as Her4 and *lfng* expression. Simultaneously, Notch signaling enhances *jag1b* and, potentially in a positive feedback loop, *notch3* expression. Together, NICD and Her6 activity may shift the network from a Delta-Notch1 lateral inhibition mode towards a Jagged-Notch3 lateral induction mode (Petrovic et al., 2014; Bocci et al., 2020), enabling stable maintenance of coherent NSC populations in NPZs. While *jagged* genes are expressed during embryogenesis (www.zfin.org) (Thisse et al., 2004; Yeo and Chitnis, 2007), read count levels in larval and juvenile scRNA-seq experiments have been too low to assign expression to clusters. We observed the regulation of *jag1b* and *jag2b* by NICD-OE at 2 dpf (Fig. S13A,D), therefore Jagged-mediated lateral induction likely emerges during larval stages. Based on the model (Fig. 5B), Delta-Notch1 and Jagged-Notch3 signaling may coexist in the ventricular layer, with higher Jagged-Notch3 activity in patches of Her6 expression. Our pseudotime analysis indicates that NPCs end *her6* expression when exiting the ventricular layer, but continue to express *delta* genes at subventricular positions, and thus may contribute Notch signaling activity to ventricular layer NSCs. *her6* and *her4* could therefore be coexpressed in NSCs, as proposed by the scRNA-seq data (Fig. 2). To determine whether in the ventricular layer *her6* is indeed co-expressed with *her4*, as indicator of Notch signaling, we analyzed their expression by HCR at 3 dpf (Fig. S14A-E). We indeed found *her4* co-expression in essentially all ventricular *her6* expression zones, indicating coexistence of Her6 and Delta-Notch-Her4 activities. To determine whether Her6 lateral induction domains may already be established at 3 dpf, we analyzed co-expression of *her6* and *jag1b* by HCR (Fig. S14F-J). While most forebrain ventricular zones have only low level dispersed *jag1b* expression, we detected *her6* and enhanced *jag1b* co-expression in the mid-hindbrain boundary (Fig. S14F) and in hindbrain proliferation zones including the rhombencephalic ventricular midline (Fig. S14H,I) and a cerebellar proliferation zone (Fig. S14J). Therefore, coherent NPZs expressing Her6 and employing Notch lateral induction may already exist at 3 dpf.

### In situ expression patterns in the adult NSC niche are consistent with a role of Her6 in locally shifting Notch signaling modes

We next investigated whether components of this regulatory network are expressed in adult NSC zones in patterns consistent with Her6 locally generating regions, in which lateral induction may prevail. At 3 months post fertilization (mpf) *jag1b* is expressed in qRGs (Morizet et al., 2024) (Fig. S5E). Therefore, we investigated the ventricular wall at the midline of the rostral telencephalon of 4 mpf adult zebrafish brains (Fig. 6). *her6*, *notch1a*, *notch3*, *lfng* and *jag1b* are expressed broadly along the midline ventricle, whereas *dla* and *dlc* show more restricted expression towards the rostral end of the telencephalic pallial ventricle (Fig. 6A1, B1, C1). We focused on this rostral region for a quantitative comparison of expression patterns (Fig. 6A-C, S15 and Methods). For each gene analyzed, transcript densities were determined based on HCR *in situ* signals within a 12 µm wide stripe lateral to the ventricular wall. Along the dorsoventral axis we defined areas based on measured *her6* transcript density: *her6*-low expressing zones below the 33^rd^ percentile and *her6*-high zones above the 67^th^ percentile of *her6* HCR signal density (Figs. S15 and 6A1, B1, C1 red and blue bars at right). Transcript densities were plotted separately for *her6-*low (dashed lines) and *her6*-high (solid lines) zones (Fig. 6A2, B2, C2).

**Figure 6.**
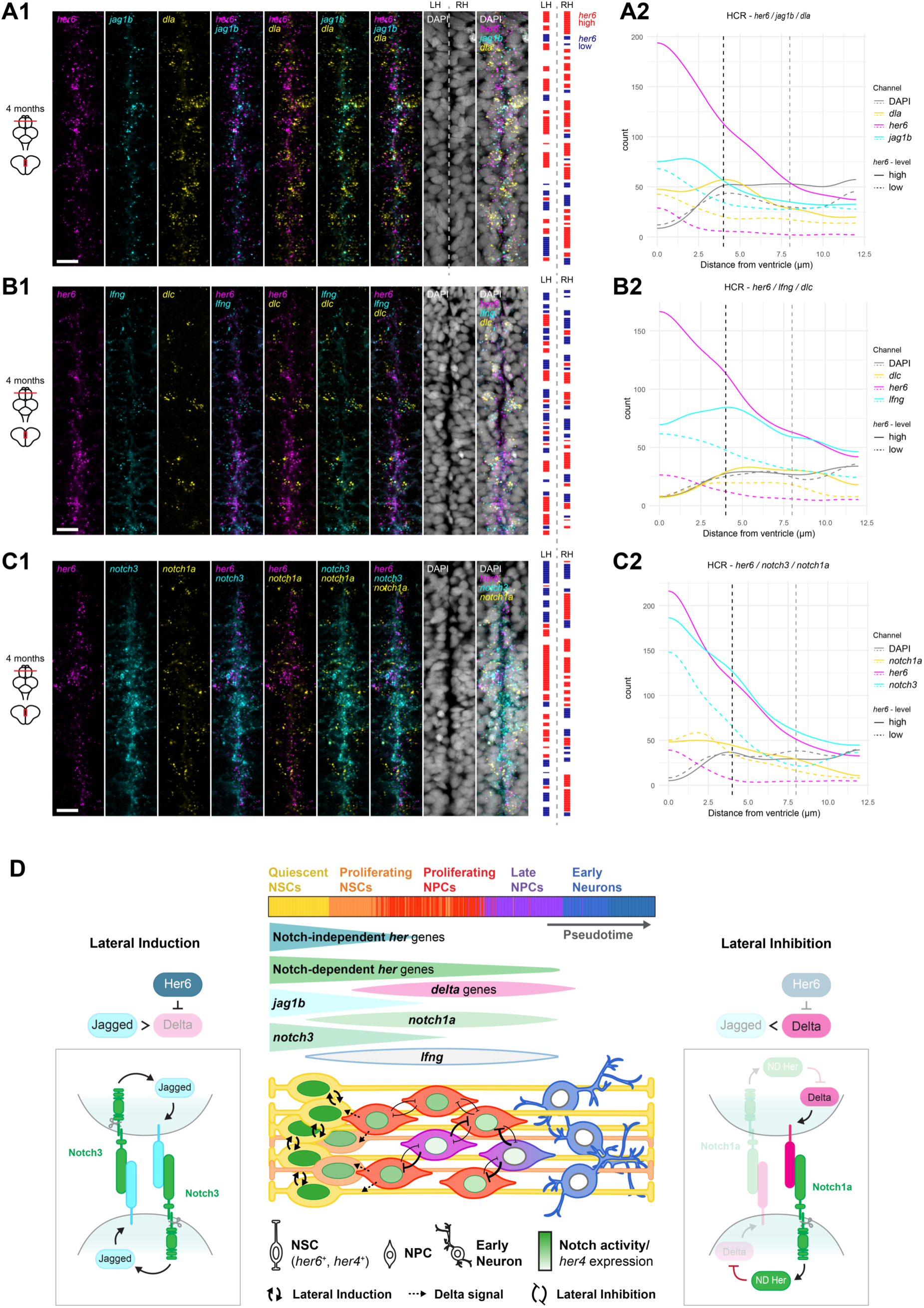
Expression of *her6* at the adult telencephalic ventricular wall in comparison to regulators of neurogenesis supports a role in control of Notch signaling modes (A1, B1, C1) In situ expression analysis (HCR) of *her6* in correlation to key components of Notch signaling involved in lateral inhibition: (**A**) *dla* and *jag1b,* (**B**) *dlc* and *lfng*, (**C**) *notch1a* and *notch3*. Images show frontal confocal sections of the dorso-rostral adult (4 mpf) telencephalon as indicate in the small brain schemes at left. The images show a 24 µm wide region centered on the midline, dorsal at top. Nuclei are stained by DAPI. **(A2, B2, C2)** Quantification of gene expression levels relative to the position of the midline (0 on x-axis) in *her6*-high (solid lines) and *her6*-low (dashed lines), for details see **Fig. S15** and Methods. Dashed vertical lines provide an estimation of the diameter of a single cell. **(D)** Schematic model of neurogenesis highlighting the activity of Her6 in control of lateral induction (left) and lateral inhibition (right) in a simplified anatomical model of zebrafish NPZs (bottom center, apical side towards ventricle at left). At top, pseudotime lineage progression (Fig. 3) is indicated aligned to the apical-basal organization of the anatomical model. Below the pseudotime is a graphical representation of temporal expression profiles of selected neurogenesis genes, correlating with neurogenesis stages in the anatomical model. Her6 represses *delta* in NSCs, promoting Notch3 activity via a Jag1b-mediated positive feedback loop, and resulting in lateral induction and maintenance of coherent fields of ventricular NSCs. As Her6 declines when NSCs progress to NPCs, *delta*- and *lfng*-driven lateral inhibition by Notch1a signaling dominates, enabling interspersed progression of neurogenesis. The anatomical model indicates that Her6-mediated lateral induction may mostly act between NSCs at the ventricular surface, while Delta signals from NPCs mediate lateral inhibition in NPCs, but at the same time activate Notch targets in NSCs through Notch3 activation. Negative regulation of *lfng* by Her6 restricts efficient Delta-Notch1 signaling to NPCs. For details see discussion.

In *her6*-high zones *her6*, *notch3* and *jag1b* transcript density peaked at the midline with a gradual decrease at increasing distance from the ventricle. In contrast *lfng*, *notch1a*, *dla* and *dlc* did not peak at the ventricular surface, but rather about one cell diameter (∼ 4 µm) lateral to the ventricular surface (**Fig. 6A1-C1**). Comparing *her6-*high and -low regions, *notch1a* spatial expression did not differ. In contrast, *lfng, dla, notch3* and *jag1b* transcripts were enriched in the *her6*-high regions. However, while *notch3* and *jag1b* HCR signal density were peaking at the ventricular surface, highest *lfng* and *dla* densities were shifted about a cell diameter away from the ventricle. Therefore, we hypothesize that in *her6*-high zones, at the ventricular surface Jagged-Notch3 signaling prevails, while Delta-Notch1 signaling and Lfng may be most active in the second cell layer away from the ventricle. Indeed, the observed higher absolute levels of *dla* and *dlc* in *her6*-high regions are consistent with a model in which Her6 promotes Delta signaling in adjacent NPCs by suppressing *delta* genes in the ventricular NSC layer. The observed transcript distributions support a model in which a mosaic pattern of multicellular patches of two types is generated, first *her6*-high ventricular patches in which lateral induction prevails to maintain coherent zones of NSCs, and second *her6*-low patches of lateral inhibition which would facilitate lineage progression and delamination into the NPC layer below the ventricular wall of NSCs (Fig. 6D).

## Discussion

NPZs and stem cell niches have to accommodate vastly different requirements, shifting from massive proliferation in NPZs during brain growth to quiescence and stable maintenance of NSCs in the adult (Schmidt et al., 2013; Urban and Guillemot, 2014; Foley et al., 2024). Cell-autonomous mechanisms, including Sox2/SoxB1 transcription factors (Graham et al., 2003; Mercurio et al., 2022), maintain the NSC lineage together with juxtacrine Notch lateral inhibition signaling to repress differentiation. However, local signaling mechanisms that maintain coherent regions of NSCs in NPZs are not well understood.

Here, we analyze NSCs and neurogenesis in the larval zebrafish brain, when NPZs are highly active. We focus on *her* genes, which either depend on Notch activity to mediate lateral inhibition, or are expressed in a NI manner to cell-autonomously maintain NSCs. Our data suggest that Notch signaling, ND Her4 and NI Her6 act in a regulatory network with distinct and specific contributions to the control of the NSC lineage. Notch and Her6 differentially affect cell cycle progression and quiescence. Further, Her6 affects the NSC transcriptional network for the Notch signaling mode to shift from lateral inhibition towards lateral induction, potentially controlling both stability and size of coherent NPZs.

### Differential combinatorial expression of NI and ND *her* genes in NSC and NPC subpopulations

Our scRNA-seq pseudotime analyses in larval zebrafish revealed transcriptome states along a continuous lineage from ncNSCs to NPCs and early neurons, which appear stable throughout larval stages, and show that the major neurogenesis lineage states are already present at 3 dpf. This is in line with previous studies describing adult-type NSC and NPC populations to already exists at embryonic and larval stages (Dirian et al., 2014; Mitic et al., 2024). NSCs and NPCs continuously express NI or ND *her* genes up to onset of *elavl3/elavl4* expression in early neurons. Co-expression of NI *her9* and ND *her4* and *her15* genes (Chapouton et al., 2011), and of *her6* and *her4* genes in adult qRGs (Cosacak et al., 2019; Lange et al., 2020) has previously been reported. We observed a progressive shift from combined NI and ND *her* expression to exclusive ND *her* expression along the lineage from ncNSCs to late NPCs. In published scRNA-seq data from the mouse dentate gyrus, *Hes1* and *Hes5* are co-expressed in astrocytes and RG-like cells from early postnatal to adult stages (Fig. 2C in Hochgerner et al., 2018 (Hochgerner et al., 2018)). Thus, simultaneous contributions of NI and ND *Hes/her* genes to regulation of NSCs appear to be prominent in vertebrates.

Expression of Hes1 and Hes5 in mice (Hirata et al., 2002; Manning et al., 2019) as well as of Her6 in zebrafish (Soto et al., 2020; Doostdar et al., 2024; Sigloch et al., 2026) oscillate. In our NSC clusters, low and high *her6* expression levels do not map to distinct subclusters, but scatter among NSC populations, indicating that “high” versus “low” *her6* oscillation phases do not establish transcriptionally distinct cell identities. The strict link of neurogenesis gene expression to pseudotime lineage rather than *her6* mRNA levels suggests that transcriptome states are driven by neurogenesis progression rather than by short Her6 oscillation phases.

### Quiescence and cell cycle control

Zebrafish quiescent NSCs of RG type were initially described in the adult brain (Chapouton et al., 2010). The origins of this population can be traced back to a population of *her4+* RGs already at 2 dpf (Dirian et al., 2014), however qRG-type NSCs have so far not been described in early larval stages. Here, we identified in 3 dpf larval brains ncNSCs exhibiting a transcriptional signature consistent with a quiescent state, similar to qRG of adult zebrafish (Lange et al., 2020; Morizet et al., 2024). Therefore, we hypothesize that ncNSCs expressing both NI and ND *her* genes at 3 dpf might establish a long-term reservoir for juvenile and adult neurogenesis.

Our OE experiments suggest distinct relative contributions of Notch signaling and NI *her6* to cell cycle control in ncNSCs. Both NICD- and Her6-OE downregulate *gadd45g*, potentially preserving Cdk1/CyclinB activity and promoting cell cycle progression. However, *NICD* also induces *gadd45ba*, which inhibits and even disassembles the Cdk1/CyclinB complex, favoring cell cycle arrest over progression. Our scRNA-seq analyses reveal high *gadd45ba* expression in non-cycling cells, suggesting a link between Notch activity and quiescence in NSCs, which has also been shown for zebrafish adult NSCs (Chapouton et al., 2010). Cell cycle arrest is reinforced by Notch OE repressing *cdc25b*, a phosphatase crucial for Cdk1 activation (Lammer et al., 1998), preventing cells from reentering the cell cycle, and together indicating G2/M arrest. While G2/M arrest in quiescent cells is documented in *Drosophila* (Otsuki and Brand, 2018; Sood et al., 2022), further research is needed to explore this in vertebrates. However, NICD-OE also induces *cdkn1cb* expression, a CDK inhibitor, indicating promotion of quiescence at G1-S transition. Cdkn1c in mice maintains undifferentiated NPCs in the neocortical ventricular zone contributing to the emergence of adult NSCs (Furutachi et al., 2015). Remarkably, Cdkn1c activities in NSCs may preserve Notch signaling effects by a positive feedback mechanism, as quiescence in murine NSCs is supported by p57 (CDKN1C) activity increasing ND non-oscillatory Hey1 expression (Harada et al., 2021).

Her6-OE primarily repressed *Cdc6*, *RHOU*, *ccnd1* and *e2f1,* genes promoting G0/G1-S transition, indicating that Her6 promotes G1 arrest at multiple levels. Notable exceptions are the *gadd45gb* paralogs and *cdc6* in its secondary role, which limit G2/M progression, and whose repression by Her6 in turn suggests support of cell cycle progression at this stage. Our data suggest that Notch activity primarily restricts G2/M progression, whereas Her6 may limit G1/S transition while maintaining the transient nature of this arrest. Concordingly, the Her6 ortholog Hes1 is downregulated in G1 for cell cycle progression, while overexpression leads to retardation of the G1 phase (Shimojo et al., 2008).

Cell culture data have revealed that high and sustained Hes1 levels are important for maintaining quiescence (Maeda et al., 2023), while the reversibility of this state is ensured by Hes1 as well (Sang et al., 2008), thereby preventing premature senescence. NI expression of Hes1 in mice has been shown to be instrumental to set aside a pool of non-proliferating “NIHes1” NSCs and maintain them as later adult NSCs (Riya et al., 2026), suggesting that some NI activities of Hes1 and Her6 in control of quiescence may be conserved.

### Control of Notch lateral induction signaling mode

Here, we identified differences in NICD, Her4.1, and Her6 transcriptional targets that together reveal a mechanism that enables Her6 specifically to shift Notch signaling modes from Delta-Notch1 towards Jagged-Notch3 signaling (Fig. 5B). Our scRNA-seq analyses revealed that *her6* is predominantly expressed in ncNSCs and pNSCs, which together with genetic evidence (Sigloch et al., 2023) indicates a pivotal role in long-term maintenance of NSCs within proliferation zones (**Fig. 6D**).

Her6 as a strong repressor of proneural genes essentially blocks neurogenesis progression, which contributes to maintaining NSCs. While Her6 also efficiently inhibits Delta expression in NSCs, these cells are still exposed to Delta signals from adjacent NPCs due to the cellular organization of the NPZ (Fig. 6D): In both larval and adult zebrafish brains, NSCs form a single cell layer at the ventricular surface (Chouly and Bally-Cuif, 2024), resulting in direct contacts with adjacent subventricular cells, including NPCs. The persistent NPC-derived Delta-dependent NICD activity may induce expression of Jagged and Notch3 in Her6-positive ventricular NSCs, where *delta* expression is repressed. This is in line with our observations following NICD-OE at larval stages, and consistent with our *in situ* expression analyses in the adult pallium. Given that NICD activation of *jag1b* has been shown to likely be direct (Ortica et al., 2026) and Notch2 as well as Notch3 are potentially more efficiently activated by Jagged than by Delta (Karlstrom et al., 2002; Kuintzle et al., 2025), we suppose that this results in a shift from Delta-Notch1 to Jagged-Notch3 signaling in Her6-positive NSCs. This shift may further be enhanced by Her6 weakly repressing *lfng*, an effect that facilitates Jagged-Notch3 signaling and attenuates Delta-Notch signaling. Finally, persistent levels of Notch activation result in positive feedback through Jagged on Notch activity.

Taken together, we propose that Her6 acts as a moderator to shift Notch signaling from lateral inhibition to a distinct signaling mode that has been previously predicted and termed lateral induction (Sjoqvist and Andersson, 2019; Bocci et al., 2020; Yoshihara and Takahashi, 2023) (Figs. 5B, 6D). In this model, increasing levels of Jagged relative to Delta drive cells from lateral inhibition (binary signal-sender-receiver cell state) towards lateral induction (hybrid signal-sender-receiver cell state), with Lfng antagonizing this process (Boareto et al., 2017). While lateral induction has been implied in several aspects of neural development (Hartman et al., 2010; de Haan et al., 2024; Ortica et al., 2026), a mechanism that controls lateral induction has been unknown so far.

Lateral induction facilitates long-term maintenance of NSCs, but also spatial organization of NSCs in proliferation zones. Further, lateral induction results in Notch activation in coherent larger groups or even sheets of NSCs, while lateral inhibition causes a salt-and-pepper distribution of NSCs versus NPCs entering neurogenesis (Sjoqvist and Andersson, 2019; Bocci et al., 2020; Yoshihara and Takahashi, 2023). In the thalamic larval proliferation zone, *her6* is expressed in a single coherent cell layer of NSCs at the ventricular wall (Sigloch et al., 2023; Sigloch et al., 2026). Our analysis of the adult pallial ventricular wall reveals that *her6, jag1b* and *notch3* expressing cells form multicellular patches of NSCs consistent with the model of lateral induction. In contrast, *dla* and *lfng* expression appear only in few interspersed cells of the ventricular wall, but predominantly in the second subventricular NPC layer, consistent with the model of lateral inhibition in NPCs.

Low residual expression of *lfng* in the Jagged-Notch3 domain versus higher *lfng* in the Delta-Notch1 domain may further refine the boundary between ventricular lateral induction and subventricular lateral inhibition zones. Although Jagged preferentially binds Notch3, it can also bind Notch1 (Shimizu et al., 1999), which may cause induction in subventricular layer cells. However, Lfng introduces inhibiting marks for Jagged1 in Notch1 (Kakuda and Haltiwanger, 2017), enhancing ligand binding but preventing trans-activation. This effectively blocks the spread of Jagged-mediated activation into the subventricular layer, while simultaneously reducing the pool of free Notch1-receptors and thereby further depleting Notch activity (Kuintzle et al., 2025). This creates an asymmetric interaction between neighboring Jagged and Delta domains: Jagged-positive ventricular cells receive Lfng-enhanced Delta-Notch signals, while in Delta-positive subventricular cells Lfng effectively shields the lateral inhibition domain from Jagged-induced Notch signal. Although a strict separation between ventricular layer lateral induction and subventricular lateral inhibition appears conceptually intuitive, the presence of *delta*- and *lfng*-expressing cells at the ventricular surface within *her6*-low zones in our analysis of adult ventricular NSCs suggests a more complex pattern, with multicellular *her6*-high patches of lateral induction in NSCs, and intermingled *her6*-low cells potentially representing intermediate NSC/NPC states

It emerges, both based on scRNA-seq and overexpression data, that NI versus ND *her* genes, Jagged versus Delta, and lateral induction versus inhibition are not strictly alternate signaling states, but more likely represent regulatory system states existing in parallel in individual cells along the ncNSC to NPC lineage progression. Indeed, our sorted Her6-mNG cells are enriched both for the Notch1 target *her4.1* and the Notch3 targets *hey1* and *jag1b* (Than-Trong et al., 2018), revealing strong Notch-signaling input into *her6* expressing NSCs.

Our scRNA-seq analyses further demonstrated that NI and ND Her activities exist in parallel in NSCs and NPCs, with the balance shifted towards NI *her* in ncNSCs and towards ND *her* in NPCs. Similarly, scRNA-seq data from the adult telencephalon (Morizet et al., 2024) also reveal both *dla* and *jag1b* in NSCs, where *jag1b* prevails in qRGs/ncNSCs and *dla* with other *delta* genes in pNSCs and NPCs (**Fig. S5**). Therefore, lateral inhibition and lateral induction may only represent extreme states within a continuum of Notch signaling activities. This dynamic network property of Notch signaling systems may accommodate for highly stable ncNSCs, but at the same time provide an exit route for lineage transitions, similar to NSC dynamics by oscillations of Her6 (Soto et al., 2020; Doostdar et al., 2024; Sigloch et al., 2026) and HES1 (Hirata et al., 2002).

Finally, Her6 may help to balance cell-autonomous versus non-autonomous mechanisms in NSC maintenance: The NSC lineage is considered to be predominantly determined by expression of Sox2/SoxB1 and other transcription factors, which primarily act autonomously, but are also regulated by complex signaling input, including regulation of Sox2 expression by Notch-RBPJk signaling (Ehm et al., 2010). Lateral induction provides a non-autonomous mechanism that enables proliferation zones to scale their ncNSC/qRG patches at the ventricular margin in accordance with spatial constraints imposed by brain growth. However, also other mechanisms such as Notch mechanosensing (Suarez Rodriguez et al., 2023) or cellular protrusions may contribute to patterning NSC clusters in the ventricular zone (Hawley et al., 2025).

### Conclusions

Notch signaling in brain development has mostly been studied in the context of lateral inhibition mechanisms in maintaining stemness and controlling lineage decisions. However, Notch also contributes to tissue self-organization through positive feedback mechanisms, including lateral induction, that involve differential Notch modification by Fringe and distinct ligand activities. Classical examples are boundary formation in the *Drosophila* wing margin by Fringe, Delta and Serrate (Panin et al., 1997), and size control of cellular fields in the vertebrate inner ear sensory epithelium, establishing the size of prosensory domains (Daudet and Lewis, 2005). Our data now suggest a novel mechanism in which NSC lineage-based NI Her6 in ventricular layer NSCs, in concert with Delta-Notch signaling also from subventricular layer NPCs, together modulate Notch1 and Notch3, as well as Delta and Jagged activity, to favor lateral induction. At the same time, both Her6 and Notch control the cell cycle independently but in a concerted fashion. We think that these mechanisms accomplish several feats: stabilize ncNSC populations, and maintain and size coherent NPZs during morphogenesis in larval as well as juvenile growth phases. While Notch signaling is intrinsically juxtacrine, lateral induction may in effect relay signals across cellular fields for size control in the NPZ. Teleosts may exploit these mechanisms to enable coherent growth of the brain over long larval and juvenile time periods, while mammals in the adult brain may have switched to more local NSC niche configurations mainly to replenish neural cells as needed.

## Experimental Procedures

### Zebrafish strains and maintenance

Zebrafish larvae of the AB/TL strain were obtained through natural breeding, kept under standard conditions and staged according to Kimmel et al., 1995 (Kimmel et al., 1995). The following transgenic lines were used: *her6:her6-mNeonGreen*^m1366^ (Sigloch et al., 2026), *Tg(hsp70l:her4.1-FLAG)^m1541^* (brief here: *hsp70l:her4.1-FLAG)* and *Tg(hsp70l:her6-FLAG)^m1492^* (brief here: *hsp70l:her6-FLAG*)(Sigloch et al., 2023), *Tg(-1.5hsp70l:GAL4)^kca4^*(brief here: *hsp70l:Gal4)* and Tg(5xUAS-E1B:6xMYC-notch1a)*^kca3^*(brief here: *UAS:myc-NICD)* (Scheer and Campos-Ortega, 1999).

### Single-cell dissociation and FACS sorting

All progeny of heterozygous *her6:her6-mNeonGreen*^m1366^ zebrafish were dissociated at 3 dpf to single cells using a protocol modified from Manoli & Driever, 2012 (Manoli and Driever, 2012) and Samsa et al., 2016 (Samsa et al., 2016) Larvae were anesthetized with 0.02% Tricaine (Sigma-Aldrich) in E3 medium (5 mM NaCl, 0.17 mM KCl, 0.33 mM CaCl_2_, 0.33 mM Mg(SO_4_)_2_) and treated 15 min with Pronase E (Sigma-Aldrich, 1mg/ml in E3) for dechorionation. Larvae were washed with E3 + Tricaine and transferred to Deyolking Buffer (55 mM NaCl, 1.8 mM KCl, 1.25 mM NaHCO_3_ in H_2_O) on ice. Deyolking was performed manually, larvae centrifuged (1 min, 270 g, 4 °C) and re-suspended for dissociation of cells in 10x TrypLE™ (10x, Gibco™, Thermo Scientific). Larvae were incubated for 20 min, and TrypLE™ digest blocked with 1x PBS (137 mM NaCl, 2.7 mM KCl, 10 mM Na_2_HPO_4_, 1.8 mM KH_2_PO_4_, Carl Roth) containing 2% Fetal Bovine Serum (Standard Quality, PPA Laboratories). The cell suspension was centrifuged (7 min, 300 g, 4°C), re-suspended, and incubated in FACSmax™ (Amsbio) two times for 30-50 min on ice, respectively. After dissociation, cells were filtered through 40 µm and 30 µm cell strainers (CellTrics® 30 µm, Sysmex) and 2 µl/ml DAPI (20 µg/ml, Sigma-Aldrich) added as viability/death indicator. Single cells were isolated und sorted by FACS (Fig. 1A). For bulk RNA-seq 4000 cells were sorted per sample, each for high, low, or no mNeonGreen fluorescence and in duplicates, respectively (FACS plot, Fig. 1A). In addition, for scRNA-seq 768 cells with high and 384 cells with low mNeonGreen fluorescence were sorted in 384 well plates.

### RNA amplification, library preparation and RNA-seq of FACS sorted cells

Cells isolated by FACS were sorted into Lysis Buffer (Herman et al., 2018) covered with mineral oil, centrifuged (3000 g, 10 min, 4 °C) and stored at −80 °C. The CEL-Seq2 protocol and library preparation were performed exactly as described in Herman et al. (Herman et al., 2018) based on the original protocol of Hashimshony et al. (Hashimshony et al., 2016). Cells for bulk RNA-seq were sorted in 12 µl Lysis Buffer and 25 µl mineral oil per sample and protocols adjusted to the volume. After the CEL-Seq2 protocol, two scRNA-seq samples of plate 2 (batch 6 and 7, with each 24 cells pooled) were mixed accidentally. As reverse transcription with barcoded primers was performed before, counts could be re-assigned after sequencing (see below). Libraries were sequenced on an Illumina HiSeq 3000 system in high-output run mode at a depth of approximately 260K reads per cell for scRNA-seq and approximately 50M reads per sample for bulk RNA-seq.

### Quantification of transcript abundance of bulk and scRNA-seq data from FACS sorted cells

Raw RNA-seq data were processed using the European Galaxy Server (Galaxy, 2024). UMI barcodes were extracted using *UMI-tools extract* (Smith et al., 2017) and reads from bulk RNA-seq trimmed according to sequencing quality (QS20) using *Trim Galore!*. For quality control *FastQC* was used and results summarized with *MultiQC* (Ewels et al., 2016). Reads were mapped to danRer10 using *RNA STAR* (Dobin et al., 2013) and Ensembl annotation GRCz10.91, with mNeonGreen sequence and annotation added, respectively. Using *Filter BAM datasets* (Barnett et al., 2011) scRNA-seq data were filtered, not allowing for multiple mapping and >2 mismatches. Gene expression was quantified for bulk and scRNA-seq data, respectively, with *featureCounts* (Liao et al., 2014). UMIs of scRNA-seq data were quantified using *UMI-tools count* (Smith et al., 2017), and bulk and scRNA-seq datasets concatenated with *Column Join*. Mixed scRNA-seq batches were rectified using *Text reformatting* and *Cross-contamination Barcode Filter* (Sagar et al., 2018). The resulting matrices were used as input for downstream data analyses in R.

### Clustering of scRNA-seq data from Her6-mNG positive FACS sorted cells

The scRNA-seq data were analyzed using *RaceID3* (Herman et al., 2018) and *VarID* (Gruen, 2020). Cells were filtered for a minimum of 1500 total transcripts, and genes with at least 3 transcript counts in 3 cells, resulting in 882 high-quality cells. Initial clustering was performed using *RaceID3* with 3 outlier genes required for outlier detection. Initial clustering revealed cells without transcript counts for *her6* and *mNeonGreen* from plates with higher fluorescence. Cluster inspection further confirmed a contamination of autofluorescent cells from the erythrocyte lineage in batches with higher fluorescence, detected by expression of hemoglobins and *alas2* (Ransom et al., 1996; Hu et al., 2014). Besides that, cells did not cluster differentially for cells FACS sorted for higher or lower mNeonGreen fluorescence, therefore “low” and “high” fluorescent cells were not handled separately in further analyses. Erythrocytes (351 cells of the *her6* and *mNeonGreen* double negative *RaceID* final clusters 1 and 10) and cells expressing any marker for eye (retina and pigment epithelium: *otx5, prdm1a, crx, nr2e3, ndrg1b, gngt2a, nme2a, rcvrn2, rpe65a* or *rlbp1a*), pigment cells (*dct, tyrp1a*, *tyrp1b* or *gch2*), neural crest (*twist1a*, *twist1b, snai1a*, *snai2, prrx1a*, *prrx1b* or *hand2*) or placodal cells (*epcam* or *cldn7b*) were excluded from the dataset for further scRNA-seq analysis. This filtering identified 146 high-quality *her6:her6-mNeonGreen* positive brain cells, for which clustering was performed with *VarID*. For clustering, we decided to use a set of 3644 genes differentially expressed (p ≤ 0.05) in our bulked mNeonGreen FACS-positive versus negative RNA-seq data (Figs. S1C, E), to counteract high variability and relatively low cell numbers, and to improve cluster discrimination.

Using marker genes for stemness (*sox2*, *sox3*, *id1*, *mdka*) and proliferation (*mcm5, mcm7*, *pcna*, *mki67*, *ccnd1*) we identified two distinct clusters of proliferating NSCs (pNSCs) and non-cycling NSCs (ncNSCs), along with a cluster of early neurons (*elavl3, elavl4*, *stmn1b*) and a non-neural cluster containing mesenchymal and muscle cells (*reck*, *lama4*, *thbs4b*, *emp2*; Fig. 1C, D). Expression of both, NI *her6* and ND *her4* genes, is highest in ncNSCs. In these cells, also *id1* and *mdka* are highly expressed, both shown to promote quiescence of RG/NSCs in adult zebrafish (Zhang et al., 2020; Lübke et al., 2022). From the literature (Lange et al., 2020) we obtained a list of marker genes enriched in quiescent and proliferating RG-like cells in adult zebrafish. To investigate whether the non-cycling Her6-mNeonGreen positive NSCs have a quiescent RG-like character, we clustered all cells from NSC clusters mNG#1 and 4 using this list of RG markers. Hierarchical clustering revealed that our ncNSCs express high levels of quiescent RG markers, while our pNSCs express proliferating RG markers (Fig. 1E).

### Analysis of bulk RNA-seq data from Her6-mNG-positive and -negative FACS sorted cells

Analyses for differential expression (DE) between bulk RNA-seq samples were performed using *EdgeR* (Chen et al., 2016). 377 genes of the erythrocyte lineage and differentially expressed (p ≤ 0.01, FC > 2) in final clusters 1 and 10 from initial *RaceID* scRNA-seq analysis of Her6-mNeonGreen positive cells were excluded (see Methods section above).

### Subsetting and clustering of scRNA-seq data from Raj et al. (2020)

Publicly available scRNA-seq data for 3, 5, 8 and 15 dpf zebrafish larval heads (3 dpf) or brains (5, 8, 15 dpf) from Raj et al. (Raj et al., 2020) were downloaded from GEO data set GSE158142. Seurat objects were updated from v2.X to v3.X using *UpdateSeuratObject* and raw counts extracted using *GetAssayData* from the *SeuratObject* Package (Hao et al., 2024). Count matrices were filtered, keeping only cells with raw count ≥1 for any of the genes *her6, her9*, *her4.1-4.5, her12, her2, her15.1-15.2, her8a, her8.2, hes6, sox2,* or *sox3* (“HER/SOX” cell set). All cells expressing marker genes for the myeloid lineage (*alas2, cahz, lyz, mpx, cpa5, spi1b, gata1a or ikzf1),* eye (*otx5, prdm1a, crx, nr2e3, ndrg1b, gngt2a, nme2a, rcvrn2, rpe65a* or *rlbp1a*), pigment (*dct, tyrp1a*, *tyrp1b* or *gch2*), neural crest (*twist1a*, *twist1b, snai1a*, *snai2, prrx1a*, *prrx1b* or *hand2*) or placodal cells (*epcam* or *cldn7b*) were removed (resulting set referred to as “EPNcPM-filtered”), as well as cells with less than 1500 total transcripts, and genes detected with less than 3 transcript counts in 3 or less cells only. The 3 dpf subsetted dataset (“HER/SOX-3dpf”) was analyzed with *VarID* using the following parameters: *knn* = 20, *nb* = 8. GO enrichment analysis of all HER/SOX-3dpf cluster defining genes (p < 0.05) revealed non-neural clusters (#5, 10, 3, 13, 12, 11, 18, 16) which were, together with an outlier cluster (#17), not considered in further analyses (Fig. S2 and grey clusters in Fig. 2C). EPNcPM-filtered HER/SOX count matrices of 3, 5, 8 and 15 dpf larval heads and brains were concatenated and analyzed with *VarID* using the following parameters: *knn* = 10, *no_cores* = 1. Default parameters were used if not indicated otherwise.

### Re-clustering of NSC and NPC clusters from HER/SOX-3dpf clustering

As stem and progenitor cell markers did not show sharp expression boundaries between clusters of pNSCs and pNPCs (Fig. 2D), we confirmed the subdivision by comparison of differentially expressed (DE) genes: Cells of the NSC clusters #14 and #15 exhibit high levels of NSC and RG markers, compared to cells of the NPC clusters #1, 8 and 9 (Fig. S3A, B).

We performed re-clustering of cells from the NSC clusters (#14 and 15; Fig. S3C) and NPC clusters (#1, 8 and 9; Fig. S3D) from HER/SOX-3dpf clustering using the same analysis pipeline with default parameters, respectively. For computation of the UMAP of NSCs the parameters *spread* = 1.8 and *n_neighbors* = 20 were used. For computation of the tSNE map of NPCs *perplexity* = 49 was used.

Re-clustering of the NSC clusters clearly distinguished non-cycling NSCs (#14) from pNSCs (#15; Fig. S3C). Cells in both clusters express the stem cell markers *sox2*, *sox3*, *id1, mdka* and *mdkb* (Fig. 2D, G), though at different levels, with higher expression in ncNSCs (#14), where cell cycle genes are downregulated. Further, genes enriched in adult quiescent RG cells (*ptn, mfge8a, cx43*) are upregulated in non-cycling NSCs (#14; Figs. 2D, G and S3C). In contrast, cells of the pNSC cluster (#15) exhibit elevated expression of early and late cell cycle genes. For re-clustering of the NPC clusters see methods section “Pseudotime Analysis” below.

### Pseudotime analysis

Pseudotime analysis was performed with *VarID* and *FateID* (Herman et al., 2018) on the HER/SOX-3dpf filtered dataset following *VarID* analysis. The trajectory, used for inspection of pseudo-temporal gene expression, was predicted based on the transition probabilities between clusters (Fig. 3A). As the two clusters of late progenitors (#8 and 9) did not have a major transition probability link (compare Fig. 3A) we confirmed that these clusters do not represent two distinct progenitor populations of different lineages: Cells of the HER/SOX-3dpf NPC clusters (#1, 8 and 9) are intermingled when re-clustered and cells from all NPC clusters are represented in populations with early cell cycle genes (G1/S), late cell cycle (G1/S) and late progenitors with *elavl3* expression (Fig. S3D). Therefore, we concluded that differential mapping in the initial clustering (Fig. 2C) was based on anatomical origin of the cells rather than different NPC lineages. Pseudotime order along a trajectory was therefore extracted for cells of HER/SOX-3dpf clusters #14, 15, 1, 9, 4, 2, 7 and 6 (Fig. 3A-C). To infer models of pseudo-temporal expression profiles self-organizing maps (SOMs) were used with the following parameters: *nb* = 50, *alpha* = 1, *corthr* = 0.95, *minsom* = 15.

### Marker genes and cluster annotation

Anatomical or cell type specific marker genes were used based on expression reported in the literature and previous scRNA-seq studies (Dulken et al., 2017; Cosacak et al., 2019; Farnsworth et al., 2020; Lange et al., 2020; Raj et al., 2020; Howard et al., 2021).

Clusters were assigned based on expression of pre-known marker genes or GO annotations (see below), and compared using information on expression from the ZFIN database (Howe et al., 2013) and the *Daniocell* online resource (v1.1, https://daniocell.nichd.nih.gov/index.html) (Farrell et al., 2018; Sur et al., 2023).

### Gene Ontology (GO) library and functional analyses

We build an extended GO library by supplementing the GO annotations for zebrafish from Ensembl GRCz10.91 with the GO annotations for mouse genes which have identified homologs in zebrafish. For mouse GO terms, we used the homology information and GO annotations of mouse markers, retrieved from the Mouse Genome Database (MGD, Mouse Genome Informatics, The Jackson Laboratory, Bar Harbor, Maine, http://www.informatics.jax.org, downloaded April 2020).

GO analyses were performed with *clusterProfiler* (Xu et al., 2024) and *enrichplot* utilizing the extended GO library. Genes with any associated GO term matching the terms “neuro”, “neural”, “nerv” or “brain” were included for identification of neuronal HER/SOX-3dpf clusters (Fig. S2B).

### Analysis of correlated gene expression using pseudo-bulking

We aimed to identify genes based on whether their expression level may be correlated or anti-correlated in relation to the mRNA levels of NI and ND *her* genes. To reduce noise and variability intrinsic to scRNA-seq data, we grouped cells into pseudo-bulk samples by assigning cells into pseudo-cells sorted by their respective expression levels separately for *her6*, *her4.1*, or *her15.1* Normalized counts for cells in clusters #14, 15, 1, 8 and 9 from clustering of the HER/SOX-3dpf EPNcPM-filtered dataset were extracted. Cells were sorted in three separate series in decreasing order by expression of *her6*, *her4.1* or *her15.1*, respectively, and in each series subsetted into groups of 12 cells (Fig. S8A). Raw counts of each group of 12 cells were merged. The resulting count matrices with pseudo-bulk samples were normalized using *VarID*. All genes with at least 1 transcript count in 1 pseudo-bulk sample were included, and distance matrices used for determining k nearest neighbors. Correlation of expression levels of each gene with expression of *her6*, *her4.1* or *her15.1*, respectively, across each series was tested using Pearson’s correlation coefficient. Both *her6* and *her4.1* show highly correlated expression with multiple stem cell and quiescence markers (Fig. S8B1, B2). In addition, several ribosomal genes are expressed anti-correlated to *her6* and *her4.1*. Together this indicates that high levels of both, NI *her6* and ND *her4.1* expression, correlate with regulation of quiescence, rather than activation of NSCs, as genes of ribosomal biogenesis have been shown to be upregulated in pNSCs (Llorens-Bobadilla et al., 2015; Dulken et al., 2017).

### Clustering of scRNA-seq data from Morizet et al., 2024

Replicates 1 and 3 from publicly available scRNA-seq data from 12 weeks adult zebrafish telencephali from Morizet et al. (Morizet et al., 2024) were downloaded from GEO: GSE225863. Datasets were analyzed using the standard workflow from the *Seurat* Package. Four principal components were used for dimensional reduction. Clusters were annotated based on marker genes and expression plotted using the *FeaturePlotFromList* function from the original publication with modifications (Fig. S5).

### Heat-shock treatment and sample collection for bulk RNA-seq time series

We used heat-shock-driven overexpression (OE) of *her6-FLAG (Tg(hsp70l:her6-FLAG)^m1492^,* and *her4.1-FLAG* (*Tg(hsp70l:her4.1-FLAG)^m1541^* (Sigloch et al., 2023) in 2 dpf zebrafish larvae. For Notch activity, we used heat-shock Gal4 driven overexpression of *notch1a-ICD* [*Tg(-1.5hsp70l:Gal4)^kca4^*and *Tg(5xUAS-E1B:6xMYC-notch1a)^kca3^* (Scheer and Campos-Ortega, 1999)]. For the overexpression experiments, *hsp70l:her4.1-FLAG* and *hsp70l:her6-FLAG* were out-crossed with *AB/TL*, while *hsp70l:Gal4* was crossed with *UAS:myc-NICD* zebrafish. Their progeny was incubated at 28.5°C until heat-shocked at 2 dpf in 50 ml Falcon tubes using E3 prewarmed to 39°C, and 15 min incubation in a 39°C water bath. After the heat-shock, larvae were returned to petri dishes and incubated at 28.5°C until sample collection. 30 heads were collected for each genotype and for each of the timepoints before (0 h) and after heat-shock (0.5, 1, 1.5, 2, 2.5, 3, 4, and 6 h). Heads were dissected by cutting from posterior of the hindbrain to the pericardial space excluding the yolk mass, and preserved in RNAlater Solution (Ambion) until RNA isolation. Samples were obtained in biological duplicates on separate days.

For the different genotypes, sample collection was as follows. *hsp70l:her6-FLAG* and *hsp70l:her4.1-FLAG* larvae were identified based on GFP expression from the *cmlc:EGFP* transgene marker in the heart. Control larvae devoid of *hsp70l:her6-FLAG* and *hsp70l:her4.1-FLAG* were identified based on absence of GFP expression, and collected separately for each transgene. Subsequently, both control samples were used as biological duplicate controls for the *her6*- and *her4.1*-overexpression analyses. For *hsp70l:Gal4*; *UAS:myc-NICD* larvae, which cannot be identified based on fluorescent transgene markers, heads were cut as described above and stored individually in 96-well plates containing RNAlater solution. Trunks were placed in corresponding 96-well positions, and used for genotyping by PCR for *gal4* and *UAS:myc-NICD*. 30 double PCR-positive larval heads were pooled for each time point. Control larvae for the *hsp70l:Gal4*; *UAS:myc-NICD* experiment were *hsp70l:Gal4* PCR-positive but *UAS:myc-NICD* PCR-negative larvae, in order to avoid detecting potential Gal4 effects on transcription.

### Genotyping of hsp70l:Gal4 and UAS:myc-NICD

To identify the *hsp70l:Gal4* or *UAS:myc-NICD* transgenes, the following primers were used:

Gal4_fwd: GATGAAGCTACTGTCTTCTATCGAACAAGC

Gal4_rev: TGAAGCCAATCTATCTGTGACGGC

UAS:myc-NICD-P009.F: CATCGCGTCTCAGCCTCAC (Scheer and Campos-Ortega, 1999)

UAS:myc-NICD-P010.R: CGGAATCGTTTATTGGTGTCG (Scheer and Campos-Ortega, 1999)

PCR results in fragments of 340 bp for *hsp70l:Gal4* and 450 bp for *UAS:myc-NICD*.

### RNA isolation, library preparation and RNA-seq of bulk samples from heat-shock time series

Heads from heat-shocked samples were dissociated and their RNA isolated using the RNAeasy Mini Kit (Qiagen). Heads collected in RNAlater Solution were transferred to RLT-buffer (RNeasy Mini Kit; Qiagen) with β-mercaptoethanol (100:1) and homogenized manually with gauge needles (Sterican 0.60 x 30 mm) attached to 1 ml syringes (B.Braun, Melsungen, Germany). RNA isolation was performed following the manufactures’ specifications, with the suggested additional DNase digestion. Library preparation and sequencing were performed by Biomarker Technologies (BMK) GmbH (Münster, Germany) using Illumina NovaSeq technology with approximately 35M reads per sample.

### Quantification of transcript abundance of bulk RNA-seq samples from heat-shock time series

Raw bulk RNA-seq data were processed using the European Galaxy Server (Galaxy, 2024). Reads were trimmed according to sequencing quality (QS20) using *Trim Galore!*. For quality control *FastQC* was used and results summarized with *MultiQC* (Ewels et al., 2016). Reads were mapped to danRer10 using *RNA STAR* (Dobin et al., 2013) and Ensembl annotation GRCz10.91, with *her6-FLAG, her4-FLAG* and *notch1a-ICD* (NCBI Accession number: XM_068215777, region: 4499-7588) sequence and annotation added, respectively. To distinguish transgene from endogenous expression in the NICD-overexpression experiment, the intracellular sequence of the Notch1a-receptor was removed from the endogenous locus. Gene expression was quantified with *featureCounts* (Liao et al., 2014). The resulting matrices were used as input for downstream data analyses in R.

We used the FLAG-tag RNA sequence to determine overexpression separate from transcript levels of endogenous *her6* and *her4.1,* as judged from read counts of non-coding parts of their transcripts that are not included in the heat-shock constructs (Fig. 4B).

### Elimination of heat-shock and Gal4 effects on transcription from time series analyses

When we normalize the time series data to the first value before heat-shock (t=0), we observe a heat-shock effect on expression of several genes in control and experimental groups at 30 min pHS (Fig. S9). The heat-shock effect decays by 60 min pHS, and can be distinguished from Her4, Her6 and NICD overexpression effects based on the temporal profile (Fig. S10). For all analyses, we compared identical time points and calculated the fold change relative to heat-shocked non-transgenic control samples for FLAG-tagged lines, and relative to Gal4-expressing control samples for Gal4-NICD-OE, thereby removing the heat-shock and potential Gal4-dependent effects from the data.

### Analysis of bulk RNA-seq data from heat-shock driven overexpression experiments

Differential expression (DE) analyses were performed using *DESeq2* (Love et al., 2014). Samples which were processed simultaneously (heat-shock and RNA isolation) were considered as batches. The Likelihood Ratio Test (LRT) was used to account for multiple parameters: *genotype*, *timescale* and *batch*. Two models were constructed: Firstly, the full model which consists of the four parameters *genotype*, *timescale*, *batch* and the interaction of *genotype* and *timescale* (*genotype:timescale*). Secondly, the reduced model which only features *genotype*, *timescale* and *batch*. The resultant adjusted *p-*value represents whether a gene is DE at one or more time points after the first timepoint. Additionally, for genes not qualifying for the strict DESeq2 p ≤ 0.05 threshold, t-tests were performed at individual time points on the difference in expression between transgenic and control larvae as indicated by light grey boxes in heatmaps (Figs. 4C, 5A, S9, S10, S11A, S12A, C, S13A-C).

### Data visualization

Data visualization was done in R using *ggplot2*, *pheatmap* and functions from the *RaceID* and *FateID* packages with modifications. Error bars in plots from overexpression experiments represent standard error of the mean, calculated by using error propagation: 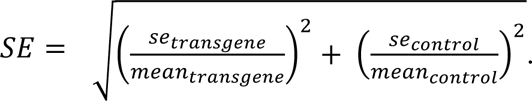. Figures were assembled and labelled in Adobe

### Sample preparation and *in situ* staining using HCR probes

The HCR probes were obtained from Molecular Instruments (Los Angeles, USA) or designed for this study. Wildtype AB/TL adult zebrafish heads were collected at 4 mpf in 4% paraformaldehyde (PFA) in PBS. After a 1-2 h fixation at 4 °C, brains were dissected from surrounding tissue in PBS and subsequently fixed in 4% PFA over night at 4 °C. Whole 3 dpf embryos were fixed under the same conditions without prior dissection. For the HCR procedure, the “HCR-RNA-FISH (v3.0) Rev. 10” protocol for whole-mount zebrafish embryos and larvae from Molecular Instruments was used, with the following modifications: After rehydration, brains were cleared in Solution 1.1 (Pende et al., 2020) for 90 min at 37°C before proceeding with proteinase K (10 µg/ml, AppliChem) in PBST (PBS with 0.1% Tween 20, AppliChem; pH 7.4) treatment for 30 min. Embryos were treated with proteinase K in the same way without prior tissue clearance. For the second overnight incubation with hairpins, the embryos were co-stained with 10 μg/ml DAPI (#D9542, Sigma-Aldrich), which was added to the amplifier solution. After the last step of the protocol, the brains and embryos were washed with 50% 5x SSCT (750 mM NaCl; 75 mM C_6_H_5_Na_3_O_7_, Carl Roth; 0.1% Tween 20; pH 7) / 50% PBST mix, followed by two 10 min PBS washes. Larvae were stored at 4°C in 80% glycerol/PBST for later imaging, while the whole brains were embedded in 3% agarose and cut in 80 µm slices from anterior to posterior with a Leica VT1000 S microtome. Brain slices were then stained with DAPI solution (10 µg/ml in PBST) for 30 minutes and stored in 80% glycerol in PBST at 4 °C.

### Imaging and analysis of HCR *in situ* stainings

Brain slices and embryos were imaged using a ZEISS LSM 880 confocal microscope with Airyscan in Fast mode. 3 dpf embryos were imaged using the LD LCI Plan-Apochromat 25x/0.8 Imm corr DIC (UV) VIS-IR objective. Brain slices were imaged using the LD LCI Plan-Apochromat 40x/1.2 Imm AutoCorr DIC M27 objective. For each brain HCR staining, either three brains with two slices each or two brains with three slices each were analysed. For analysis of the transcript distribution, the dorsal ventricular region in the anterior telencephalon was virtually straightened using the “Straighten” ImageJ plugin (Kocsis et al., 1991), and then cropped to 12 µm width from the midline ventricle into each hemisphere. To identify transcripts separately, the ImageJ function “Find Maxima” (https://imagej.net/ij/docs/guide/146-29.html) was used with default settings (Prominence > 10) and maxima counted using the ImageJ “Measure” function. Subsequent analyses were performed in R, starting with splitting the measured transcript positions to analyse each hemisphere separately. We further distinguished between *her6*-high and *her6*-low zones by dividing the images in zones along the ventricle based on transcript density at the 33^rd^ and 67^th^ percentile over all analyzed brain slices of each staining. Counts were then summed up in a kernel density estimate function in ggplot2 with a bandwidth of 1 µm for *her6*-high and -low zones respectively.

Along the medio-lateral axis, transcript distribution may be affected by mechanisms of apical mRNA localization (Kusek et al., 2012; Vessey et al., 2012), however, we did not observe strictly apically localized HCR signals at higher magnification, and conclude that changes in transcript density may mostly reflect differences in expression between the cell layers adjacent to the midline.

## Supporting information

PDF with assembled Suppementary Figures S1 to S15 and legends

## Resource Availability

Requests for further information, resources, and reagents should be directed to Wolfgang Driever.

## Ethics statement

All animal experiments were approved by the ethics committee for animal experiments at the state authority Regierungspräsidium Freiburg with the following permit numbers: Regierungspräsidium Freiburg AZ 35-9185.64/1.1, AZ 35-9185.81/G-12/40, AZ 35-9185.81/G-16/96, and AZ 35-9185.81/G-23/076.

## Acknowledgements

We thank Josip Hermann, Nina Peltokangas and Sagar, as well as the FACS and Sequencing Facility at the MPI for help with RNA amplification, library preparation and RNA sequencing. Thanks Sabine Götter for excellent fish care. The authors acknowledge the support of the Freiburg Galaxy Team: Mehmet Tekman and Prof. Rolf Backofen, Bioinformatics, University of Freiburg, Germany funded by Collaborative Research Centre 992 Medical Epigenetics (DFG grant SFB 992/1 2012) and the German Federal Ministry of Education and Research (BMBF grant 031 A538A de.NBI-RBC).

## Funding Statement

Funded by the Deutsche Forschungsgemeinschaft (DFG, German Research Foundation) under Germany’s Excellence Strategy CIBSS—EXC-2189—Project ID 390939984 (W.D., D.G.), CRC 850 Project ID 89986987 (W.D.) and GRK2344 MeInBio Project ID 322977937 (R.P., W.D., D.G.).

## Author Contributions

W.D., J.R. and R.V. conceived the study. R.V. designed and performed Her6-mNeonGreen RNA-seq experiments, bioinformatic analyses, and assembled figures. J.R. designed and performed time series experiments, RNA-seq, bioinformatic analyses and HCR-WISH, and assembled figures. J.R. created the supplemental software package for time series overexpression analysis. D.S. and J.S. contributed to time series data generation. C.S. provided the *her6:her6mNeonGreen* transgenic model. D.G. designed and supervised scRNA-seq experiments. J.R., R.V. and W.D. wrote and edited the manuscript with input from all authors, who approved the final manuscript and submission.

## Declaration of Interests

The authors declare no competing interests.

## Supplementary Information

Supplementary Figures S1-S15 are contained in the Supplementary Materials PDF. Additional supplementary data will be made available upon peer review journal publication.

## Notes

### Competing Interest Statement

The authors have declared no competing interest.

### Summary of Updates

We edited abstract, introduction, results and discussion for clarity. We exchanged all figure (1 through 6) images in the main manuscript for higher resolution images in order to generate a PDF file with sufficient detail of the figures. The original contents of the figures is unchanged. We added DOIs to the reference list.

