## Supplementary material for "Notch-independent Her6 contributes to control of neural stem cell maintenance by shifting Notch signaling from lateral inhibition towards lateral induction mode": PDF with assembled Suppementary Figures S1 to S15 and legends

**Inventory of Supplementary Figures**


**Fig. S1** Her6 positive cells feature enriched NSC markers but depleted NPC and early neuron markers

**Fig. S2** Characterization of neural and non-neural clusters of HER/SOX-3dpf cells

**Fig. S3** Differential expression of quiescence and stemness marker genes in NSC and NPC clusters

**Fig. S4** Characterization of HER/SOX clusters of the neural lineage and combinatorial expression of *her/hes* genes

**Fig. S5** Gene expression profiles of selected neurogenic, proneural, and cell cycle genes in clusters from adult telencephalic scRNA-seq analysis

**Fig. S6** Pseudo-temporal expression profiles of proneural and neurogenic genes and genes controlling cell cycle progression and arrest

**Fig. S7** All juvenile brain HER/SOX neurogenesis cell states, including non-cycling NSCs, are already present at 3 dpf

**Fig. S8** Genes with expression levels highly correlated to *her6* or *her4.1* mRNA levels in cluster maps cluster with NSC or NPC cell states rather than with *her6* or *her4.1* expression levels in individual cells

**Fig. S9** Heat-shock effect on gene expression is temporally distinct from effect of overexpression of *her4.1*, *her6* or *NICD*

**Fig. S10** Heat-shock induced overexpression of *her4.1*, *her6* or *NICD* reveals shared and distinct downstream transcriptional targets

**Fig. S11** Effects of *her4.1*, *her6* or *NICD* overexpression on cell cycle control genes

**Fig. S12** Expression levels of *her/hes/hey* and proneural genes are differentially affected by overexpression of *her4.1*, *her6* or *NICD*

**Fig. S13** Effects of *her4.1*, *her6* or *NICD* overexpression on Notch, Wnt and BMP signaling pathway components

**Fig. S14** Co-expression analysis of *her6* and *her4*, and correlation of *her6* and *jag1b* expression in 3 dpf zebrafish larvae

**Fig. S15** Quantification method of HCR in-situ imaging data of 4 mpf zebrafish telencephalons

### Supplementary Figures

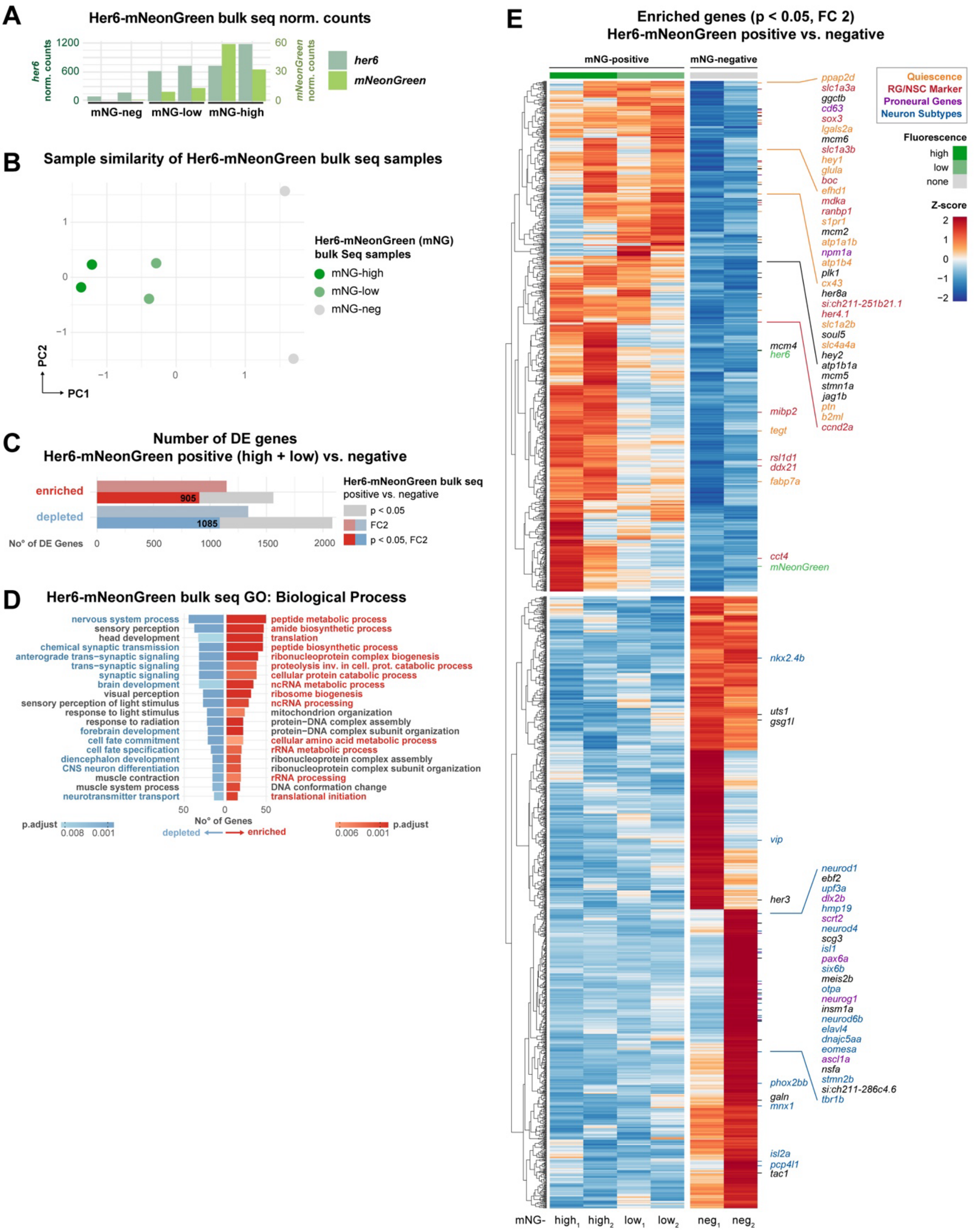


---

**Supplementary Figure S1 (Related to Figure 1B).**

**Her6 positive cells feature enriched NSC markers but depleted NPC and early neuron markers**

**(A)** Normalized *her6* and *mNeonGreen* transcript counts in bulk RNA-seq of FACS-purified 3 dpf Her6-mNG cells. Shown are biological duplicates for mNG-negative, mNG-low and mNG-high fluorescence cells.

**(B)** Sample-to-sample similarity of duplicates is shown by principal component analysis (PCA) of bulk

RNA-seq samples, showing that at the level of the whole transcriptome, low and high Her6-mNG bulk RNA-seq samples are different.

**(C)** Differential expression analysis of Her6-mNG-positive (high + low combined) vs. -negative bulk RNA-seq samples revealed expression of 905 genes enriched and of 1085 depleted genes ($p<0.05$ and FC2).

**(D)** Result of Gene Ontology (GO) analysis for significantly enriched and depleted genes ($p<0.05$, FC2) from **C**. Top 20 GO terms (p adj <0.05) are shown. GO terms associated with neural development (blue) or metabolic processes (red) are highlighted. GO analysis revealed that genes associated with neuronal differentiation are depleted in *her6* expressing cells.

**(E)** Heatmap based on Z-score of normalized expression of 905 enriched and 1085 depleted genes ($p<0.05$, FC2) in Her6-mNG positive (high and low each with replicates) and negative (neg) samples from bulk RNA-seq.

Selected NSC and neurogenesis marker genes are shown. Her6-mNG positive samples reveal depletion of genes associated with neuronal development, synaptic differentiation and cell fate commitment (**D**). Among these genes are proneural genes (*ascl1a*, *neurog1*) and genes for neuron differentiation (*neurod1*, *neurod6b*, *neurod4*, *elavl4*) and subtype specification (*otpa*, *isl1, isl2b*, *nkx2.4b*, **E**).

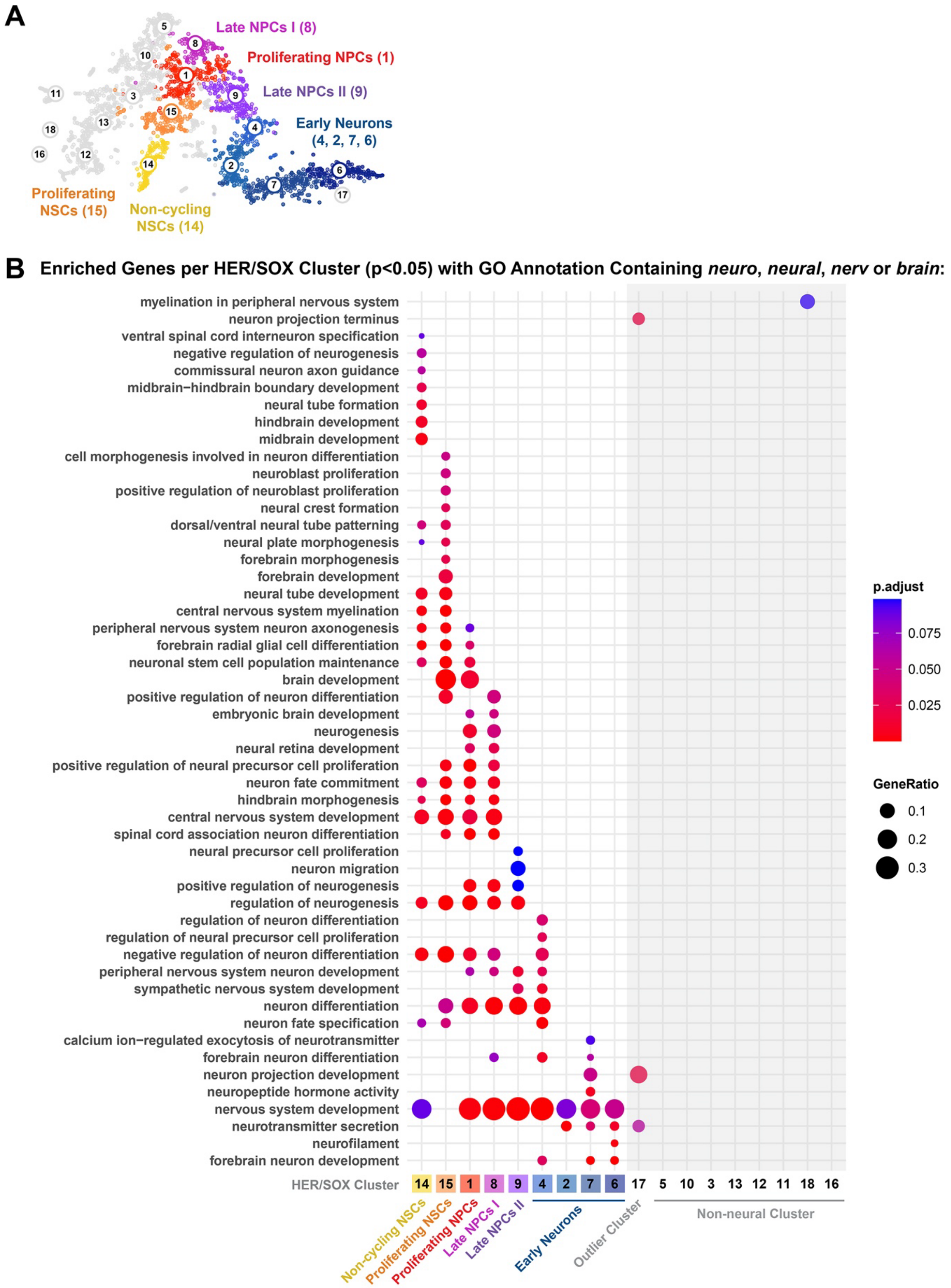


**Supplementary Figure S2 (Related to Figure 2)**

**Characterization of neural and non-neural clusters of HER/SOX-3dpf cells**

**(A)** UMAP representation of *VarID* clustering from scRNA-seq analysis of HER/SOX-3dpf cells (see **Fig. 2C**).

**(B)** Neural versus non-neural clusters of HER/SOX-3dpf cells are distinguished by GO analysis including all cluster defining genes ($p<0.05$). Only GO annotations with GO terms containing “neuro”, “neural”, “nerv” or “brain” were considered. Outlier clusters and those identified as non-neural are indicated in grey.

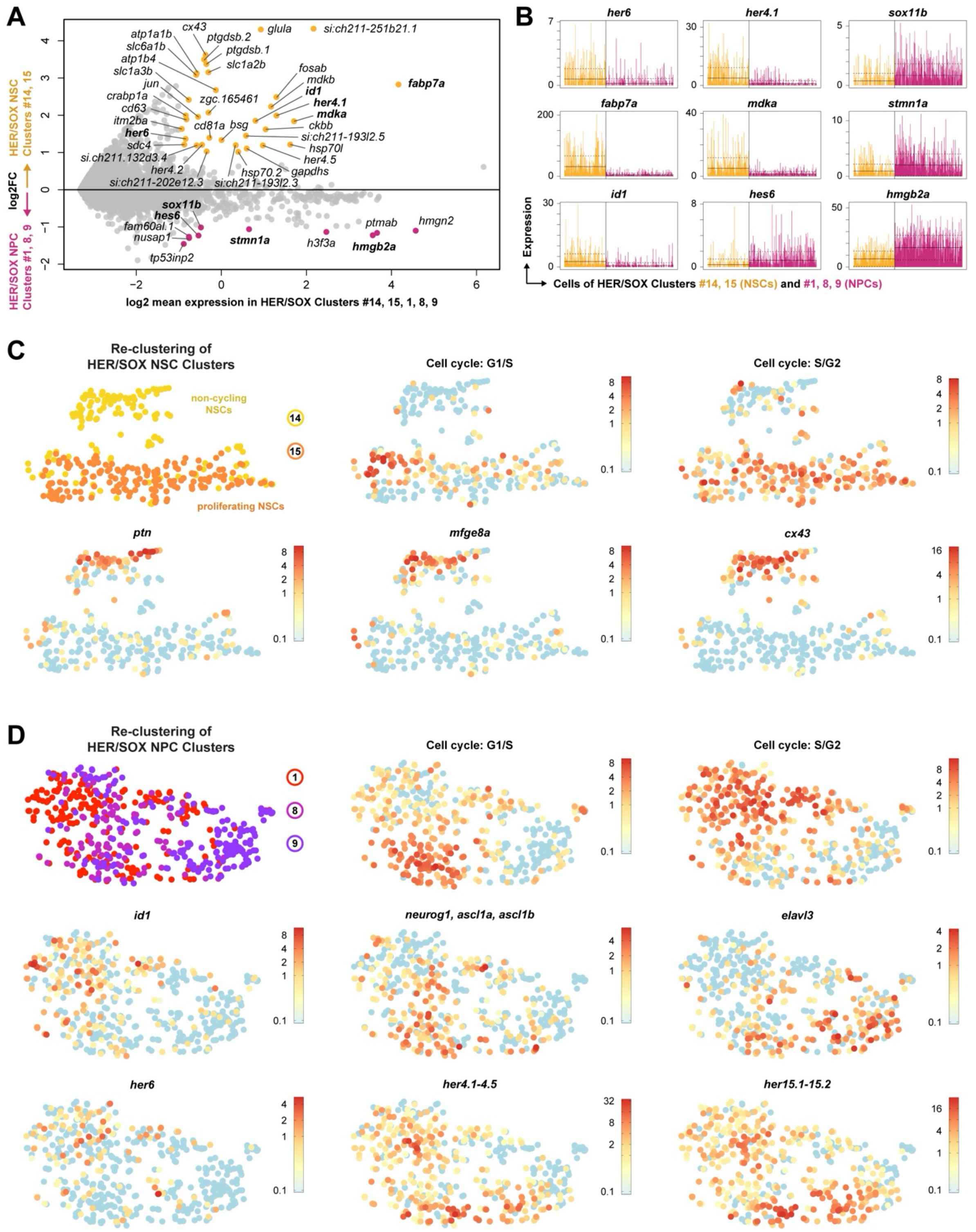


**Supplementary Figure S3 (Related to Figure 2)**

**Differential expression of quiescence and stemness marker genes in NSC and NPC clusters**

**(A)** MA plot of genes differentially expressed between HER/SOX-3dpf NSC clusters (#14 and 15) and NPC clusters (#1, 8 and 9) from **Fig. 2C**. Genes with p<0.01 and FC2 are shown.

**(B)** Barplot with expression of selected genes from **A** per cell of NSC clusters #14 and 15 (yellow) and NPC clusters #1, 8 and 9 (magenta).

**(C,D)** Reclustering of NSC and NPC clusters. Cell cycle markers for G1/S plots include *ccnd1, ccnd3*, *ccne1*, *ccne2*, *cdt1*, *mcm5*, *nus1*, *pcna*, *skp2*, *tmem2,* and for S/G2 plots include *ccna1, ccna2*, *ccnb1*, *ccnb2*, *ccnb3*, *ube2c*.

**(C)** UMAP representation of clustering from *VarID* analysis by re-clustering 309 cells from HER/SOX-3dpf NSC clusters #14 and 15 from **Fig. 2C**, with cluster annotation retained from initial clustering. Log2 normalized expression of selected genes.

**(D)** tSNE map of clustering from *VarID* analysis by re-clustering 502 cells from HER/SOX-3dpf NPC clusters #1, 8 and 9 from **Fig. 2C**, with cluster annotation retained from initial clustering. Log2 normalized expression of selected genes as indicated.

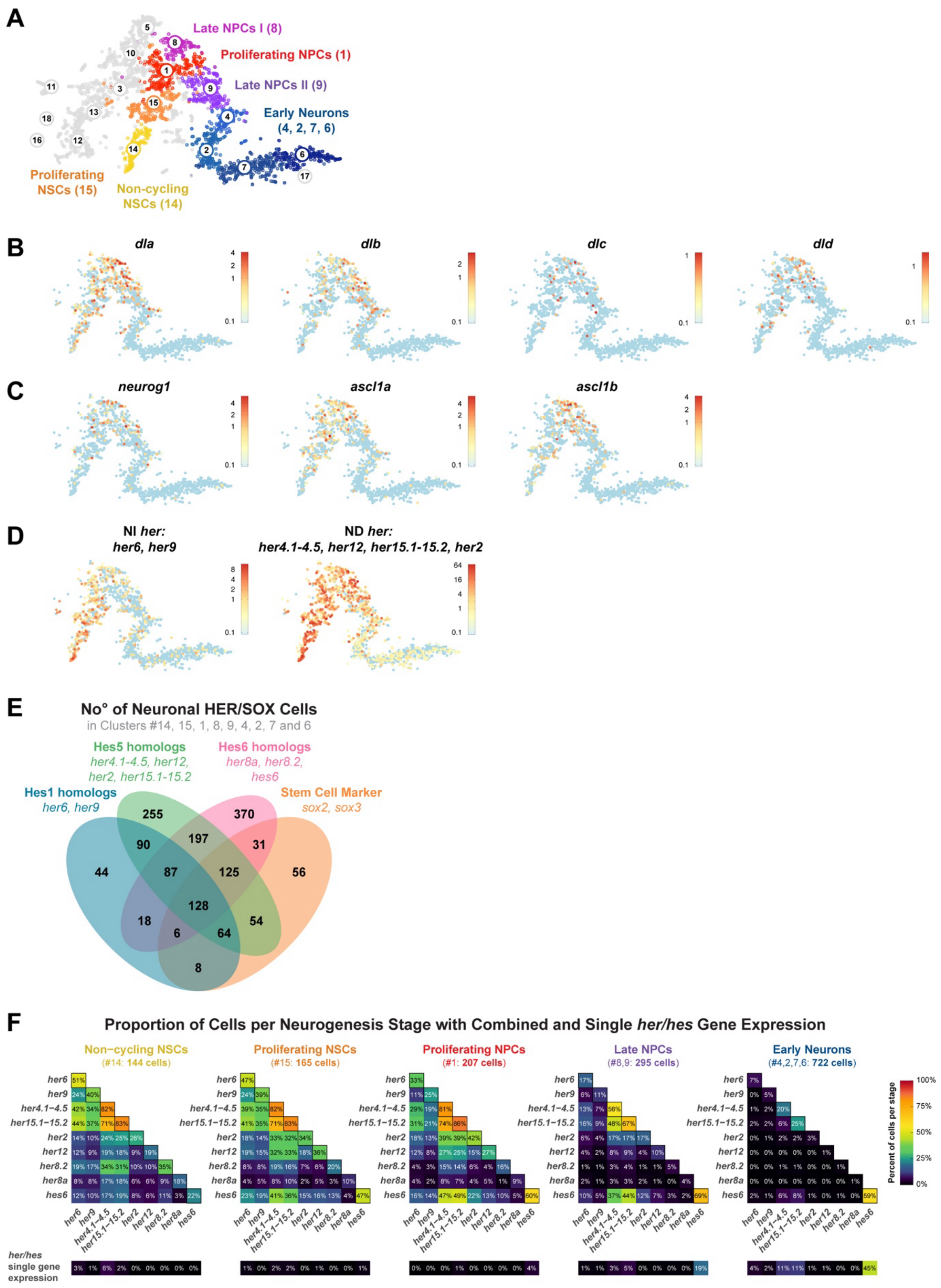


**Supplementary Figure S4 (Related to Figure 2)**

**Characterization of HER/SOX clusters of the neural lineage and combinatorial expression of *her/hes* genes**

**(A)** UMAP representation of *VarID* clustering from scRNA-seq analysis of HER/SOX-3dpf cells (see

**Fig. 2C**).

**(B-D)** Log2 normalized expression of *delta* genes (**B**), proneural genes (**C**) as well as combined NI and combined ND *her* genes (**D**) in clusters #14, 15, 1, 8, 9, 4, 2, 7 and 6 of clustering from **Fig. 2C**.

**(E)** Venn diagram illustrating distribution of individual and combined *her* class (Hes1, Hes5 or Hes6 homologs) and *sox2/sox3* gene expression detected in individual high-quality neural HER/SOX-3dpf.

**(F)** Matrices showing for each neurogenesis stage the proportion of cells with combinatorial expression of two or more *her/hes* genes (top) or exclusive expression of a single *her/hes* gene (bottom). Numbers give percentage of cells expressing a specific combination in relation to all cells analyzed at this neurogenesis stage as indicated in matrix headers, respectively. The matrices should be read as follows (example, starting at top): Non-cycling NSCs - 51% of 144 cells express *her6* alone or in combination with any other *her/hes* gene (matrix), however, only 3% of all cells express exclusively *her6* (bottom row). 24% of cells express a combination of *her6* and *her9* (and potentially any other *her/hes* gene).

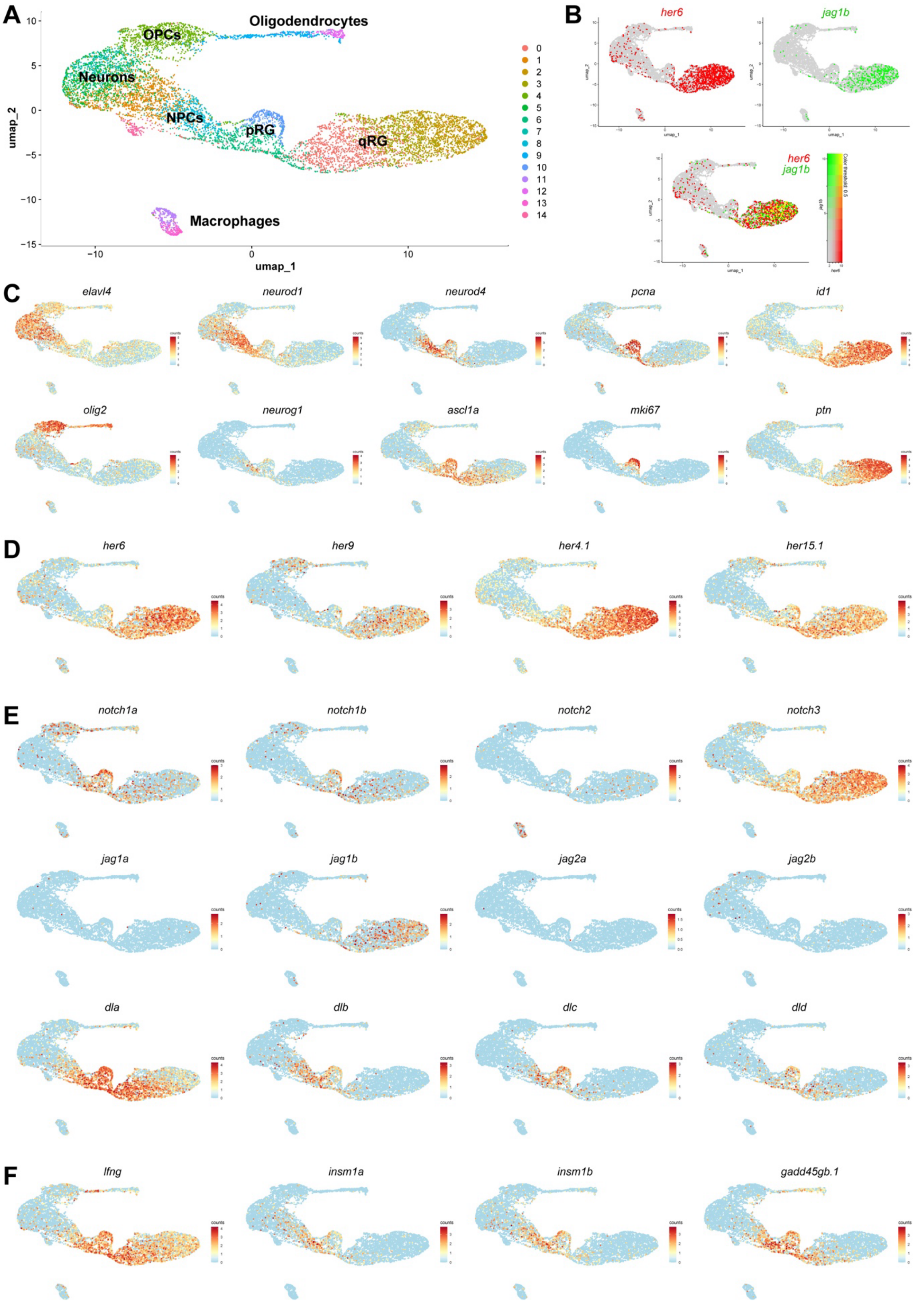


**Supplementary Figure S5 (related to Figures 2 and 6)**

**Gene expression profiles of selected neurogenic, proneural, and cell cycle genes in clusters from adult telencephalic scRNA-seq analysis**

**(A)** UMAP representation of *Seurat* clustering from analysis of publicly available scRNA-seq data from telencephalic cells from adult zebrafish from Morizet et al., 2024 (Morizet et al., 2024). OPCs, oligodendrocyte precursor cells; NPCs, neural progenitor cells; pRG, proliferating radial glia; qRG, quiescent radial glia.

**(B)** Co-expression of *her6* and *jag1b* in clustering from **A**.

**(C-F)** Expression of neurogenesis and proliferation markers (**C**), selected *her* genes (**D**), Notch receptors and ligands (**E**) and selected genes (**F**) in clustering from **A**.

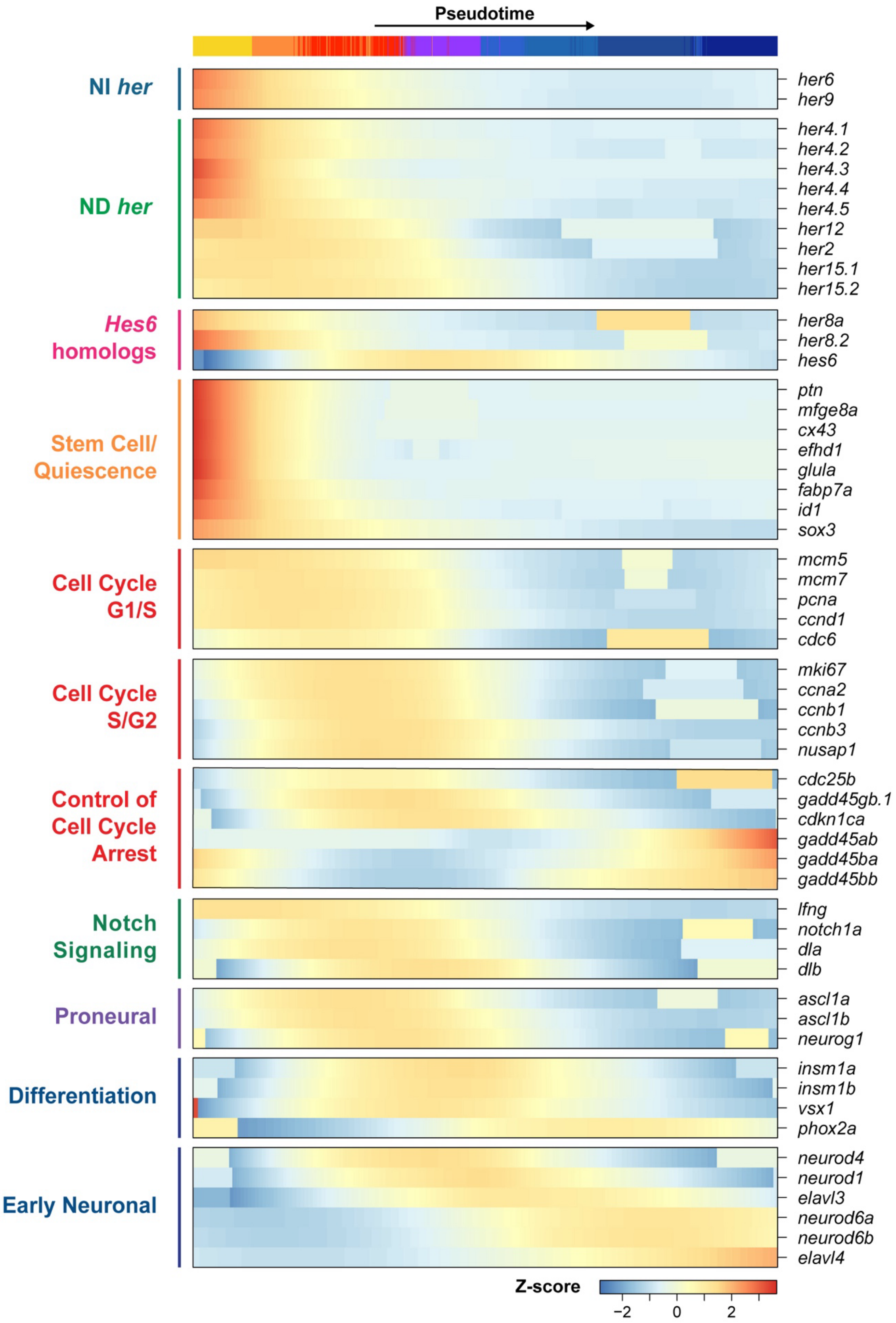


**Supplementary Figure S6 – (Related to Figure 3)**

**Pseudo-temporal expression profiles of proneural and neurogenic genes and genes controlling cell cycle progression and arrest**

Heatmap with Z-score pseudo-temporal expression profiles of selected genes for cells in pseudo-temporal order. Colors (top) indicate cluster affiliation of cells as indicated in Fig. 2.

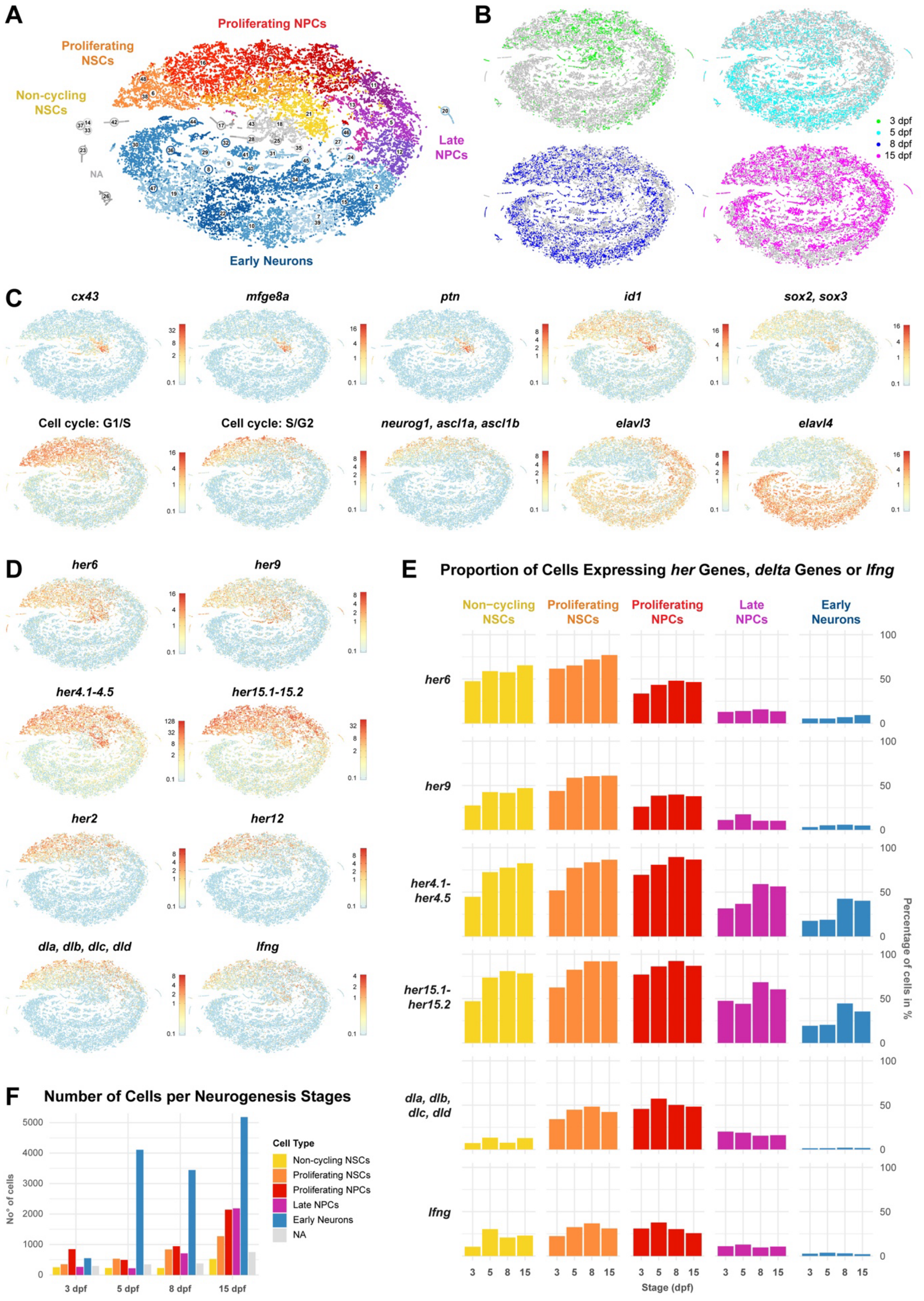


**Supplementary Figure S7 (Related to Figure 2)**

**All juvenile brain HER/SOX neurogenesis cell states, including non-cycling NSCs, are already present at 3 dpf**

**(A)** tSNE map of clustering from *VarID* analysis of 27,069 HER/SOX cells (see Methods) from publicly available scRNA-seq data from heads of 3, 5, 8 and 15 dpf larval zebrafish from Raj et al., 2020 (Raj et al., 2020). Major cell types are indicated.
**(B)** tSNE map highlighting cells of each developmental stage, respectively.
**(C)** Log2 normalized expression of selected neurogenesis and proliferation marker genes in clustering from **A**. Cell cycle markers for G1/S include *ccnd1, ccnd3*, *ccne1*, *ccne2*, *cdt1*, *mcm5*, *nus1*, *pcna*, *skp2*, *tmem2,* and for S/G2 include *ccna1, ccna2*, *ccnb1*, *ccnb2*, *ccnb3*, *ube2c*.
**(D)** Log2 normalized expression of selected *her/hes* genes and Notch signaling components in clustering from **A**.
**(E)** Proportion of cells summarized for major cell types per stage expressing *her* genes, *delta* genes or *lfng*. The data indicate that the specific expression profile for ncNSCs is already established at 3 dpf and consists at least into juvenile 15 dpf stages.
**(F)** Number of cells analyzed are summarized for major cell lineage states per developmental stage.

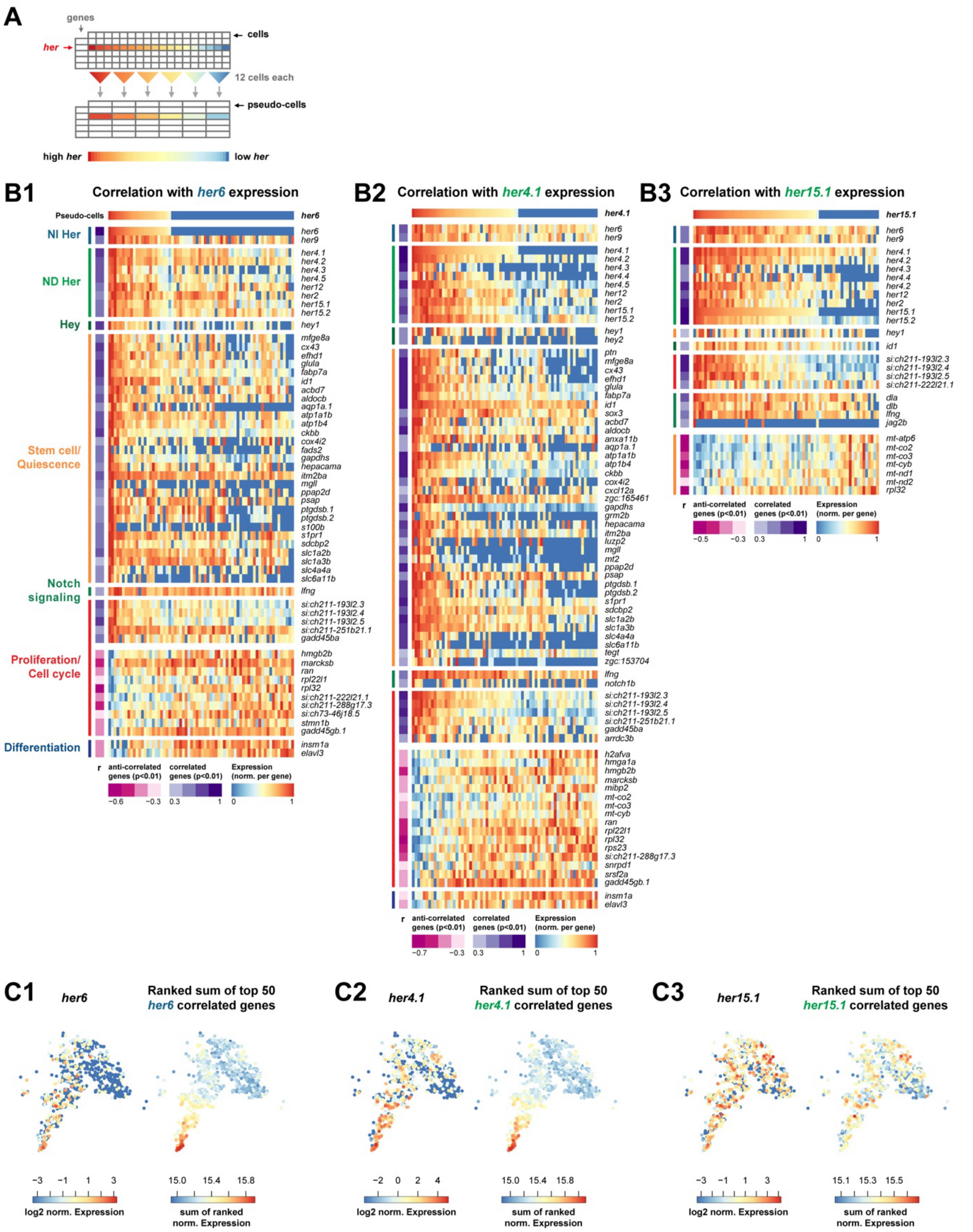


**Supplementary Figure S8 (Related to Figure 2)**

**Genes with expression levels highly correlated to *her6* or *her4.1* mRNA levels cluster with NSC or NPC cell states rather than with *her6* or *her4.1* expression in individual cells.**

**(A)** Schematic representation of pseudo-bulking, used to identify genes expressed correlated or anti-correlated to *her6*, *her4.1* and *her15.1* mRNA levels, respectively. Cells from HER/SOX-3dpf NSC and NPC clusters #14, 15, 1, 8 and 9 from **Fig. 2C** were ordered by expression level of the specific *her* gene. Based on this high to low expression sorted list, read counts of 12 consecutive cells each were merged into pseudo-cells (**see Methods**).

**(B)** Heatmap with normalized expression of selected correlated (top, purple marks) or anti-correlated (bottom, magenta marks) genes ($p<0.01$) for *her6* (**B1**), *her4.1* (**B2**) or *her15.1* (**B3**) in pseudo-cells- Expression level of each *her* genes in each pseudo cell is indicated at top. Pearson's correlation coefficient (*r*) was used to determine correlation.

**(C)** Log2 normalized expression of *her6* (**C1**), *her4.1* (**C2**) or *her15.1* (**C3**) in NSC and NPC clusters #14, 15, 1, 8 and 9 of clustering from **Fig. 2C** (**left**). Normalized counts of top 50 with correlated expression were ranked and the summed expression plotted, respectively (**right**). The expression of genes correlated to *her4.1* or *her6* expression levels maps with the NSC and NPC states, but not to the *her* levels in individual cells, indicating that the expression levels of these genes are not determined by oscillating Her expression in individual cells, but by progression of neurogenesis.

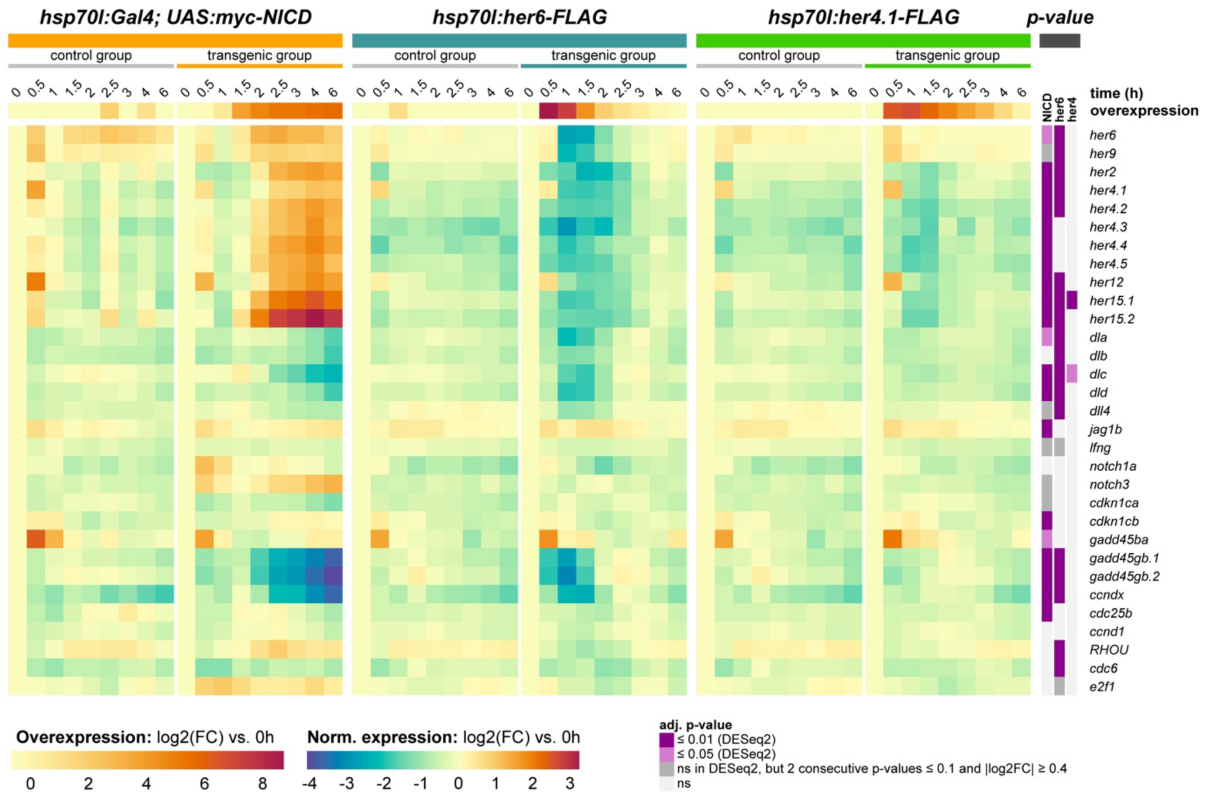


**Supplementary Figure S9 (Related to Figures 4, 5)**

**Heat-shock effect on gene expression is temporally distinct from effect of overexpression of *her4.1, her6* or *NICD***

As control for experiments in Figs. 4C and 5A, this heatmap shows log2-fold change over t=0 for expression of selected neurogenesis and cell cycle genes. Heat-shock effects on transcription are mostly limited to the first hour after heat-shock and do not interfere with time series analysis of the overexpression experiment, since (i) OE downstream genes are mostly activated only from 1 h post heat-shock onward, and (ii) the calculation of OE effects eliminated the heat-shock effect by using heat-shocked non transgenic samples as controls. The effects of ectopic Gal4 expression alone can be estimated by comparing the control group from the NICD-OE expression experiment to the control groups of Her6-OE and Her4-OE (for example, the Gal4 control group reveals that heat-shock Gal4 results in slightly increased endogenous *her6* expression).

The quality of control samples can be assessed in the first row indicating the levels of ectopically expressed *myc-NICD*, *her6-FLAG* and *her4-FLAG,* respectively. Here, it seems a small contamination of *Tg(UAS:myc-NICD)* fish was present in samples 2.5h and 4h post heat shock, while the 1h control sample for the *her*-OE experiments shows a slight Tg(hsp70l:her6_FLAG) contamination. As the control samples served as a baseline for the overexpressing samples, the major conclusions drawn from these data are not affected, besides potentially missing out on DE genes with very subtle effects only at the affected time points.

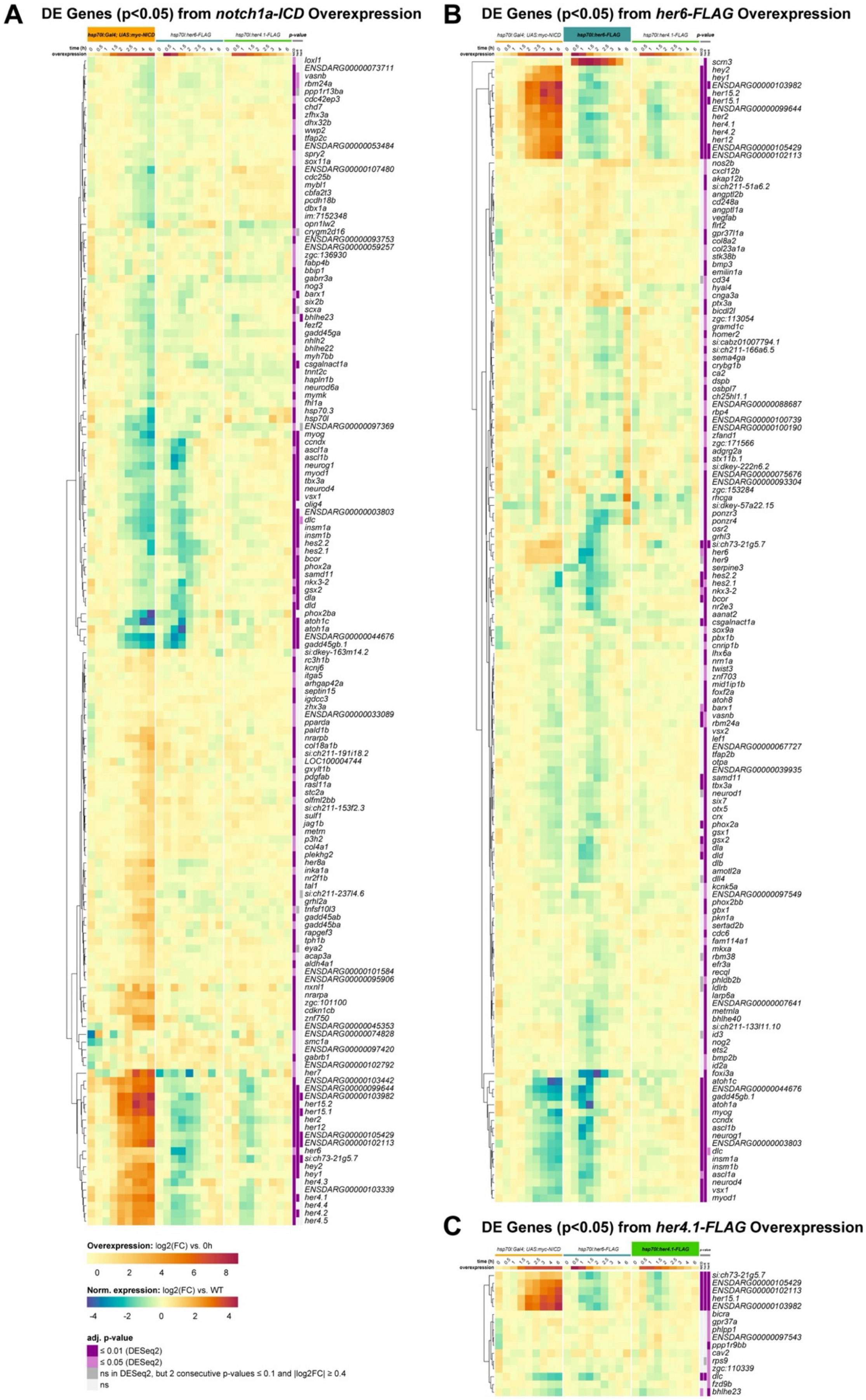


**Supplementary Figure S10 (Related to Figure 4, 5)**

**Heat-shock induced overexpression of *her4.1*, *her6* or *NICD* reveals shared and distinct downstream transcriptional targets**

Heatmaps with log2(FC) expression of all genes differentially expressed (p adj. ≤ 0.05) upon overexpression of *myc-NICD* (**A**), *her6*-FLAG (**B**), or *her4.1*-FLAG (**C**) in bulk RNA-seq time series at one or more time points after the first timepoint. First timepoint is before heat-shock.

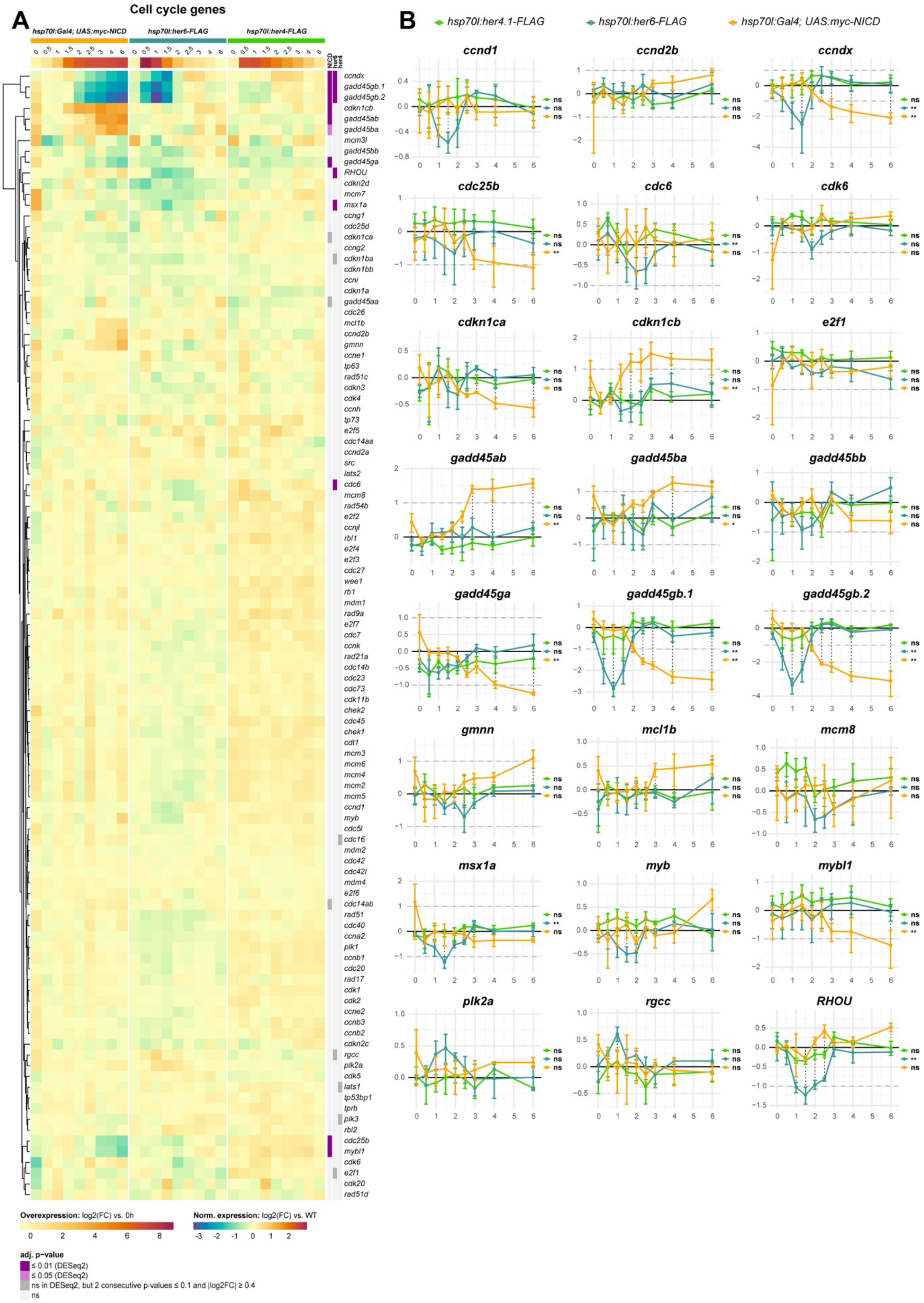


**Supplementary Figure S11 (related to Figure 4)**

**Effects of *her4.1*, *her6* or *NICD* overexpression on expression of genes with GO annotations related to cell cycle**

(**A**) Heatmaps with log2(FC) expression of selected genes based on GO annotations related to cell cycle in bulk RNA-seq time series at timepoints before and after heat shock induced overexpression of *myc-NICD*, *her6-FLAG* or *her4.1-FLAG*, respectively (**Fig. 4**). (**B**) Log2(FC) ± SEM expression of selected genes from (**A**).

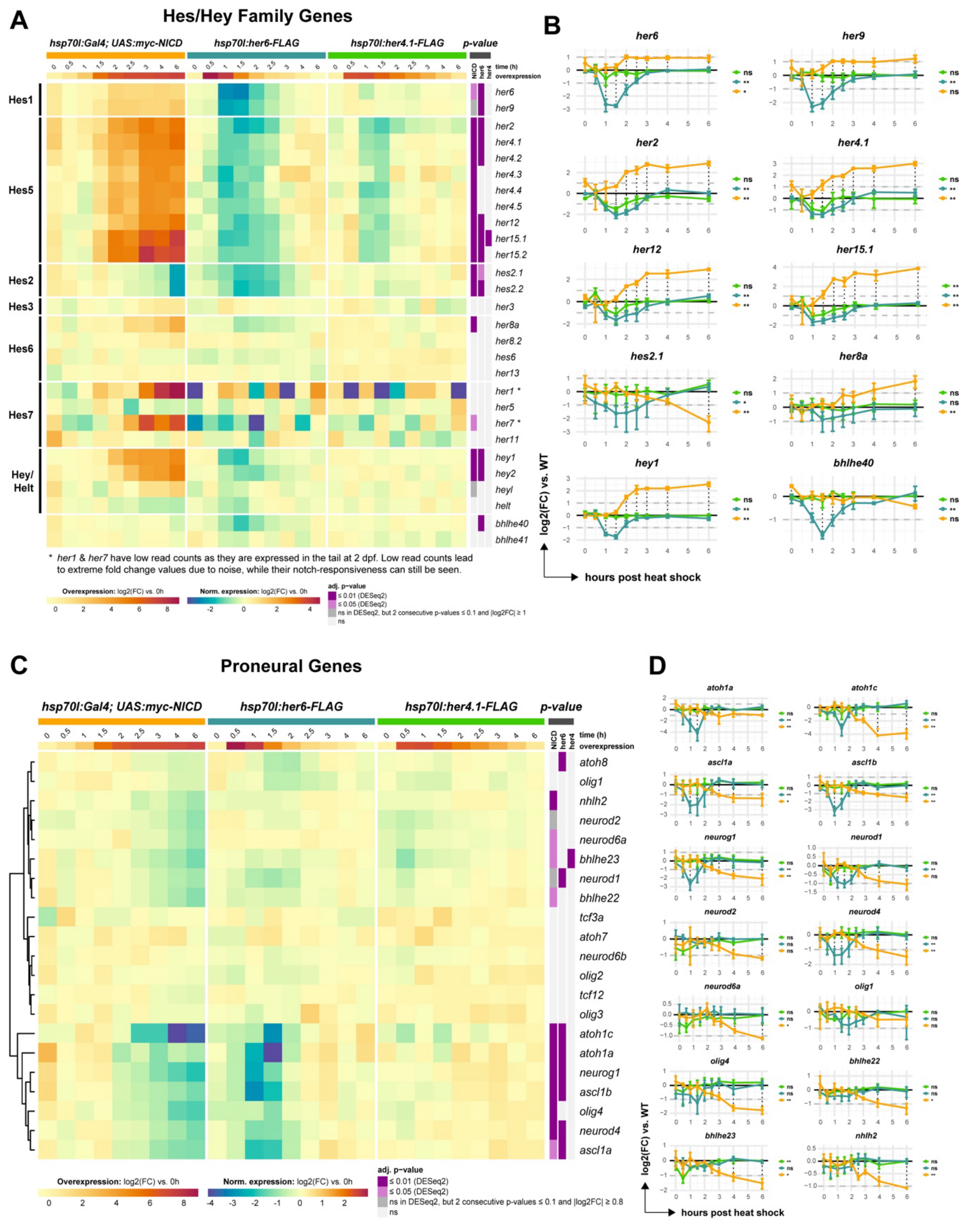


**Supplementary Figure S12 (Related to Figure 5)**

**Expression levels of *her/hes/hey* and proneural genes are differentially affected by overexpression of *her4.1*, *her6* or *NICD***

**(A)** Heatmap with log2(FC) expression of genes from the *her/hes/hey* gene family in bulk RNA-seq time series before and after heat shock induced overexpression of *myc-NICD*, *her6-FLAG* or *her4.1-FLAG*, respectively (**Fig. 4A, B**). We included *hes7* homologs, but note that *her1* and *her7* are expressed in somitogenesis but have low read counts in the brain, resulting in noisy signal.

**(B)** Log2(FC) ± SEM expression of selected genes from the *her/hes/hey* gene family.
**(C)** Heatmap with log2(FC) expression of proneural/neural bHLH genes (as compiled in (Bertrand et al., 2002)).
**(D)** Log2(FC) ± SEM expression of selected proneural genes.

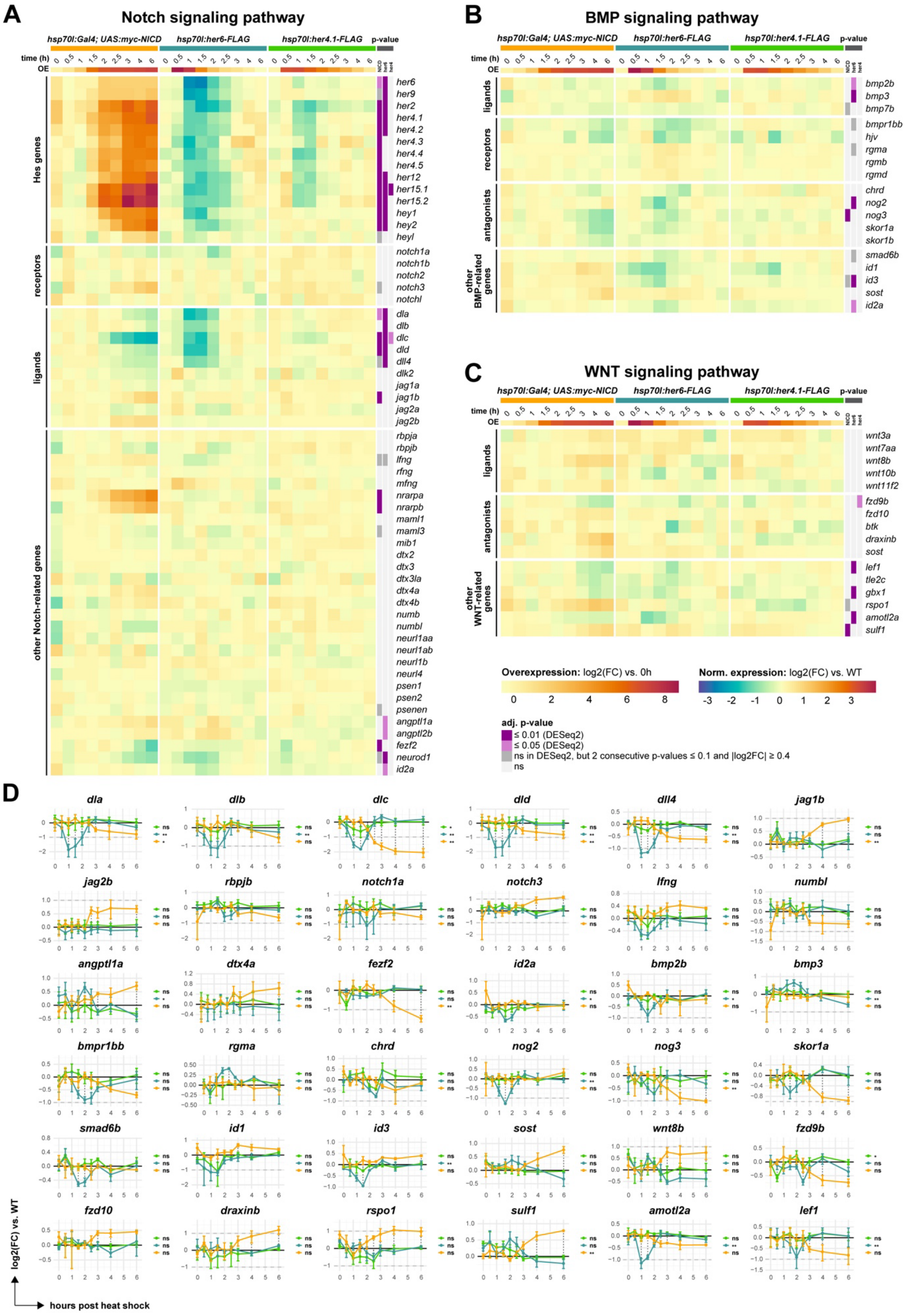


**Supplementary Figure S13 (Related to Figure 5)**

**Effects of *her4.1*, *her6* or *NICD* overexpression on Notch, Wnt and BMP signaling pathway**

**components**

Heatmaps with log2(FC) expression of manually selected genes involved in Notch (**A**), BMP (**B**) or Wnt (**C**) signaling in bulk RNA-seq time series at timepoints before and after heat shock induced overexpression of *myc-NICD*, *her6-FLAG* or *her4.1-FLAG*, respectively (**Fig. 4A, B**).

(**D**) Log2(FC) ± SEM expression of selected genes from (**A**)-(**C**).

**(B-C)** Identification of potential cross-talk of the Her6 and NICD regulatory network with other signaling pathways previously linked to stem and progenitor cell regulation. **(B)** *her6* OE repressed, but NICD OE induced expression of the BMP targets *id1 (her6*: p adj = 0.77; *NICD*: p adj = 0.91*)* and *id3 (her6*: p adj = 0.00018; *NICD*: p adj = 1*)*. *her6* OE repressed BMP signaling pathway components *nog2* (*her6*: p adj = 0.00014), *smad6b* (*her6*: p adj = 0.58, $2cFC_{p0.1} \geq 0.5$) and *id2a* (p adj = 0.042). *her6* OE and *NICD* OE both repressed *bmpr1bb* (*her6*: p adj = 0.56, $2cFC_{p0.1} \geq 0.8$; NICD: p adj = 1), *nog3* (*her6*: p adj = 0.9; *NICD*: p adj = 0.0055) and *skor1a* (*her6*: p adj = 0.32; *NICD*: p adj = 0.17). While *her6* OE induced *bmp3* (p adj = 4.4e-05), it repressed *bmp2b* (p adj = 0.042). **(C)** *her6* OE and *NICD* OE repress the Wnt signaling pathway components *amotl2a* (*her6*: p adj = 4.1e-09; *NICD*: p adj = 1), *lef1* (*her6*: p adj = 0.0046; *NICD*: p adj = 35), *fzd9b* (*her6*: p adj = 0.41; *NICD*: p adj = 0.87), *znf703* (*her6*: p adj = 0.012; *NICD*: p adj = 81), *tle2c* (*her6*: p adj = 1; *NICD*: p adj = 1) *and gbx1* (*her6*: p adj = 0.001; *NICD*: p adj = 1). In contrast, *NICD* OE enhanced *wnt8b* (p adj = 0.64), *rspo1* (p adj = 0.74, $2cFC_{p0.1} \geq 1$), *sulf1* (p adj = 0.0015) and *sost* (p adj = 0.47) expression. Examination of our observations in context of pathway function revealed both positive and negative modulation of pathway activity. Therefore, our experiments may not reveal a global interaction of Notch with BMP and WNT pathways, but may reflect diverse effects on distinct anatomical regions and cell states.

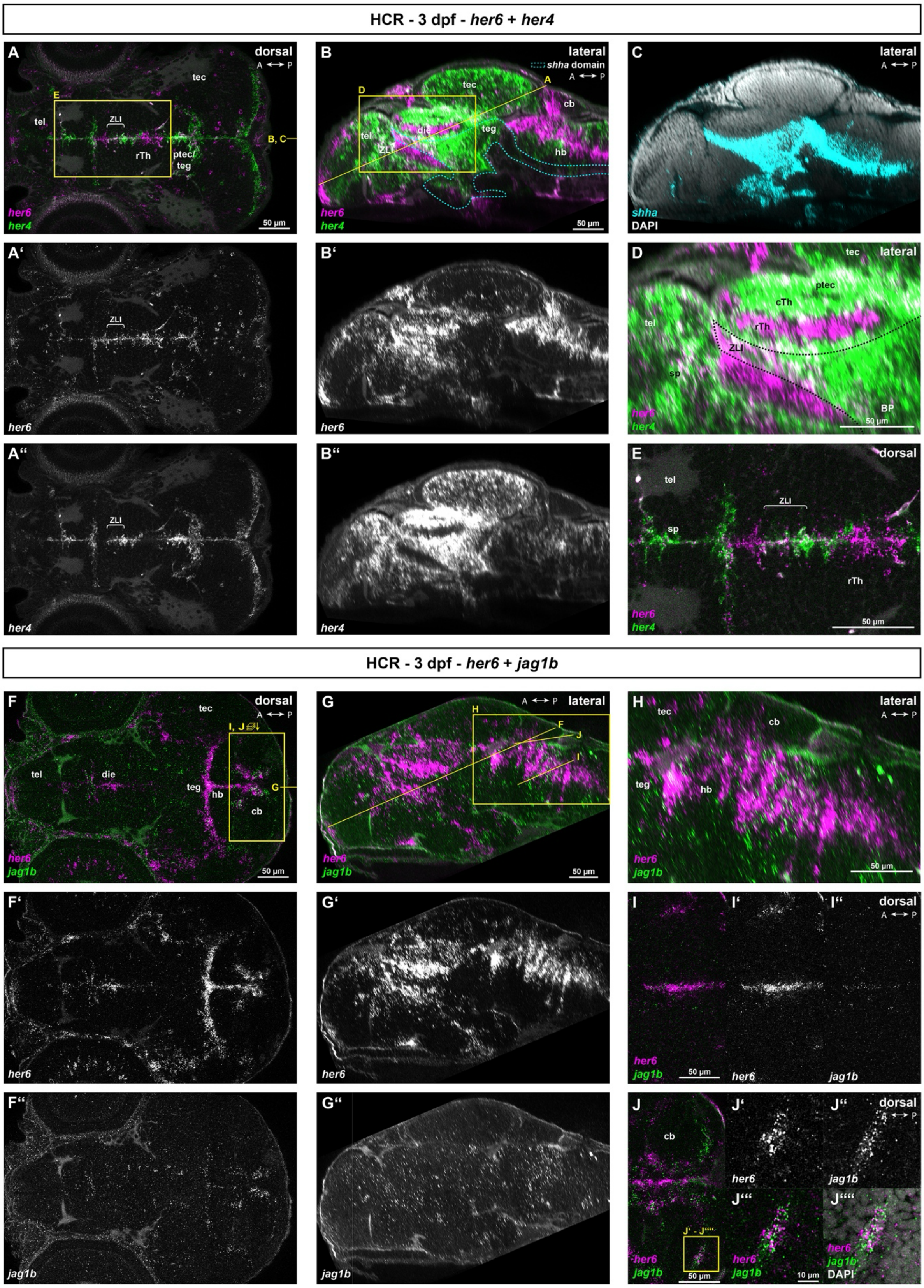


**Supplementary Figure S14 (Related to Figure 5)**

**Coexpression analysis of *her6* and *her4*, and correlation of *her6* and *jag1b* expression in 3 dpf zebrafish larvae**

**(A-J)** Coexpression analysis using HCR detection of mRNAs as indicated in whole mount 3 dpf brains.

Confocal stacks of whole heads were recorded in dorsal views (A, E, F, I, J), and sagittal views (B, C, D, G, H) generated by re-slicing of the volume. Positions of additional slices and magnified regions indicated in yellow. **(A-E)** HCR detection of *her4.1, her6,* and for anatomical orientation *shha* (C)*;* nuclei stained with DAPI. Selected channels shown as indicated. **(A, E)** Dorsal view of the fore- and midbrain, revealing that *her4* expressing cells (A'') along the midline ventricular wall also coexpress *her6* (A'), albeit generally at lower levels. **(B)** Midline sagittal reslice of (A) demonstrating coexpression of *her4* within *her6*+ domains in the telencephalon (except for the olfactory bulb), zona limitans intrathalamica, the rostral edge of the rostral thalamus, the caudal thalamus and pretectum. **(D)** Enlarged sagittal view of the telencephalic and diencephalic *her*-expression domains. **(E)** Enlarged view of the forebrain ventricular region, showing coexpression across all *her4*-expressing domains. **(F-J)** HCR detection of *her6* and *jag1b;* nuclei stained with DAPI. **(F)** Dorsal view of *her6* and *jag1b* expression domains across the fore-, mid- and hindbrain. While *her6* expression in the forebrain shows little correlation with *jag1b* transcripts, colocalization can be detected in cerebellar and rostral hindbrain proliferation zones adjacent to the tegmentum. **(G)** *jag1b* has only dispersed low level expression long the telencephalic ventricular wall, while in the hindbrain, *jag1b* expression appears enriched in *her6* domains (magnified in **H). (I)** Hindbrain ventricular layer *her6* expression domain with colocalized *jag1b* low level HCR signal **(J)**. The cerebellar ventricular zone shows strong coexpression of *her6* and *jag1b*, consistent with a potential role in establishing lateral induction domains. Abbreviations: A: anterior; cb: cerebellum; cTh: caudal thalamus; die: diencephalon; hb: hindbrain; rTh: rostral thalamus; P: posterior; ptec: pretectum; sp: subpallium; tec: tectum; teg: tegmentum; tel: telencephalon; ZLI: zona limitans intrathalamica.

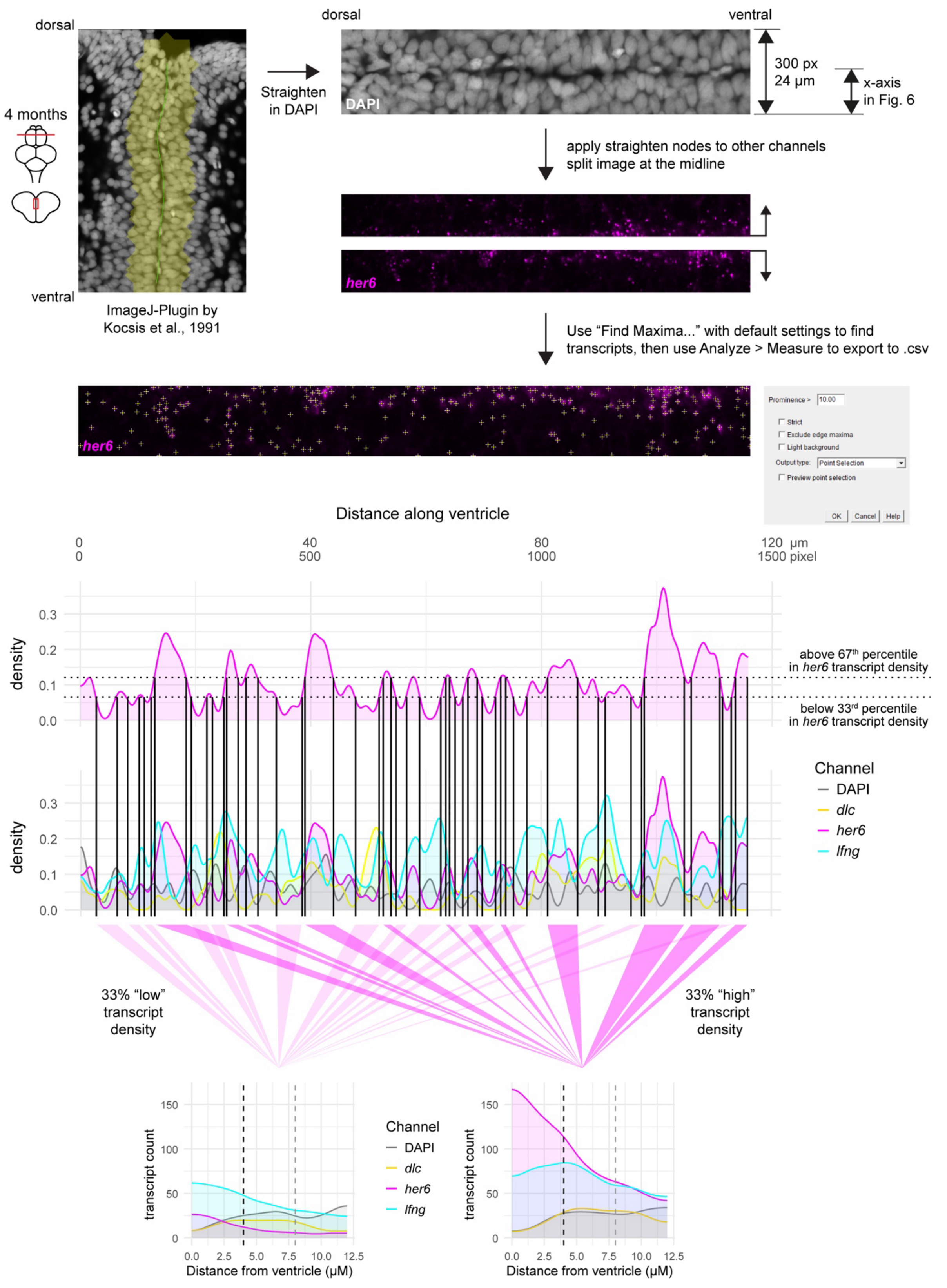


**Supplementary Figure S15** (Related to Figure 6)

**Image analysis of HCR in situ expression data in Fig. 6A-C and quantification of HCR transcript spots in *her6*-high and -low zones**

Confocal optical sections in frontal orientation were acquired at the midline from corresponding telencephalic regions of 4 mpf zebrafish brains (3 imaging planes each for 2 brains or 2 imaging plains each for 3 brain). The images were straightened along the ventricular surface using the *Straighten* plugin(Kocsis et al., 1991). Images were cropped to mediolateral dimension of 150 pixels (12 µm, about 3 cell diameters) into each hemisphere (see **Supplementary Data SD4**). *her6* transcripts were detected separately in each hemisphere using the ImageJ *Find Maxima* tool with default settings, and their positions were processed into density distributions by kernel density estimation. Density profiles from all images of a given staining were pooled to define thresholds for the upper and lower thirds of *her6* transcript density, establishing zones of *her6*-high and -low along the ventricle. Transcripts from other channels were then quantified relative to these zones and their distance from the ventricle. The example image shown here corresponds to the left hemisphere in **Fig. 6B**.
